# Parasite-driven cascades or hydra effects: susceptibility and foraging depression shape parasite-host-resource interactions

**DOI:** 10.1101/2021.05.27.445996

**Authors:** Jason C. Walsman, Alexander T. Strauss, Spencer R. Hall

## Abstract

1. When epidemics kill hosts and increase their resources, should the density of hosts decrease (with a resource increase, this constitutes a trophic cascade) or increase (a hydra effect)? Seeking answers, we integrate trait measurements, a resource-host-parasite model, and experimental epidemics with plankton. This combination reveals how a spectrum from cascades to hydra effects can arise. It reflects tension between parasite-driven mortality (a density-mediated effect) and foraging depression upon contact with parasite propagules (a trait-mediated one).
2. In the model, mortality rises when higher susceptibility to infection increases infection prevalence. Epidemics release resources while suppressing hosts (creating a cascade). In contrast, when hosts are less susceptible and parasites depress their foraging, a resource feedback can elevate host density during epidemics (creating a hydra effect), particularly at higher carrying capacity of resources. This combination elevates primary production relative to per-host consumption of resources (two key determinants of host density).
3. We test these predictions of the qualitative effects of host traits and resource carrying capacity with trait measurements and a mesocosm experiment. Trait measurements show clonal lines of zooplankton hosts differ in their foraging depression and susceptibility. We seeded resource-host-parasite mesocosms with different host genotypes and provided different nutrient supplies to test model predictions. Hydra effects and trophic cascades arose under different conditions, as predicted by the model.
4. Hence, tension between trait-mediated and density-mediated effects of parasites governs the fate of host density during epidemics – from cascades to hydra effects - via feedbacks with resources.

## 1. Introduction

Virulent parasites threaten host populations across taxa (Dobson *et al*. 2008). Parasites exact fitness costs, increasing mortality and/or reducing fecundity of infected hosts. This fitness harm can decrease host density (Daszak, Cunningham & Hyatt 2000), potentially contributing to extinction of host populations (Ebert, Lipsitch & Mangin 2000; Vredenburg *et al*. 2010). Additionally, harmful outbreaks of parasites can severely damage crops (Fry & Goodwin 1997) and livestock (Cleaveland, Laurenson & Taylor 2001; Horan & Fenichel 2007). Furthermore, epidemics that depress host density can trigger conservation crises, e.g. in mammals (Roelke-Parker *et al*. 1996), birds (Cooper *et al*. 2009), and amphibians (Vredenburg *et al*. 2010). Declines of host populations during epidemics can also indirectly release resource species consumed by hosts in parasite-driven trophic cascades (Buck & Ripple 2017). These increased resources may then increase host fitness, making the net effect of parasites on host populations more complex. These net effects matter for community composition (Wood *et al*. 2007) or biological control (Boivin, Hance & Brodeur 2012); such concerns emphasize the value of understanding and predicting whether and how strongly parasites will suppress host density and release their resources.

The strength of parasite-driven trophic cascades should depend upon the traits that control interactions among parasites, hosts, and their resources. One key trait is susceptibility of hosts to infection. Since susceptibility promotes infection of new hosts, it typically increases the proportion of hosts infected (Dwyer & Elkinton 1993; Thrall & Burdon 2000; Strauss *et al*. 2015; Strauss *et al*. 2018). Once infected, individual hosts can suffer virulent reductions of fitness such as increased mortality (Ebert, Lipsitch & Mangin 2000). Multiplied by higher prevalence, such virulence increases population-level mortality and depresses host density (Hochachka & Dhondt 2000; Hall *et al*. 2011). Therefore, higher susceptibility should lead to stronger ‘trophic cascades’, with stronger host suppression and larger resource release. In this sense, susceptibility to parasites acts like the attack rate of predators in predator-prey-resource systems: both modulate mortality of victims and should increase cascade strength (see Appendix: section 1a or Shurin & Seabloom 2005). Hence, enemy-enhanced mortality can broadly strengthen trophic cascades.

However, natural enemies can also depress their victim’s foraging rate. For example, cues from predators can reduce foraging rates of their prey, e.g., if prey reduce foraging to avoid predation (Morgan 1988; Laundré, Hernández & Ripple 2010). Similarly, the foraging rate of hosts can slow due to virulent effects of infection, behavioral response to infection, or behavioral avoidance of propagules (Raveh *et al*. 2011; Hite *et al*. 2017; Strauss *et al*. 2019; Hite, Pfenning & Cressler 2020). Foraging reduction might penalize hosts with lower nutritional intake and fecundity (Buck, Weinstein & Young 2018). Then again, hosts which eat propagules of parasites might benefit from reduced exposure via foraging reduction (Hite, Pfenning & Cressler 2020). Additionally, such foraging depression should also indirectly release resources from consumption pressure (Philpott *et al*. 2004). For hosts that slow their foraging in response to parasites, higher resource density may compensate for slower feeding. Hence, depressed foraging could impose mixed consequences for both prey and hosts: costly reduction of energy intake could be offset by released resources and lower mortality (Morgan 1988; Beckerman, Uriarte & Schmitz 1997). The net effect of foraging depression in these scenarios is unclear.

Here, we show how the interplay between mortality and foraging depression controls the strength of parasite-driven trophic cascades but can also produce hydra effects. In a hydra effect, a source of mortality (e.g., predator or parasite) leads to higher – not lower – density (e.g. of prey or hosts); one mechanism for this is via a trait-mediated indirect effect on the victim’s resources (Matsuda & Abrams 2004). Resource release can support more or fewer victims, depending on how resource production is divided among victims (Schröder, van Leeuwen & Cameron 2014). When foraging depression increases production more than per capita consumption of resources, a hydra effect arises via “prudent resource exploitation” (Abrams 2009). Alternatively, if the increase in resource consumption is stronger, then enemies drive a trophic cascade. We apply this theory to parasites that cause both mortality (a density-mediated effect) and foraging depression (a trait-mediated effect). Further, we predict and empirically test how host and resource traits govern the range from hydra effects to strong trophic cascades during epidemics.

Hence, we illustrate how susceptibility, foraging depression, and carrying capacity of the resource lead to both cascades and hydra effects in a single system by interweaving models and experiments. We measured susceptibility and foraging depression traits of several clonal genotypes of zooplankton hosts and used them to parameterize the model. The model predicts that a virulent parasite drives stronger trophic cascades as susceptibility of hosts increases (with or without foraging depression). However, hosts with lower susceptibility but strong foraging depression can experience hydra effects, especially at high carrying capacity of the resource. We tested these predictions using those same host genotypes in a mesocosom experiment. As predicted, populations of more susceptible hosts suffered larger epidemics (higher prevalence) of a fungal parasite. These larger epidemics more strongly suppressed host density and released algal resources. Further, hydra effects emerged for resistant (low susceptibility) genotypes with strong foraging depression in systems receiving high nutrients. Hence, parasites (like predators) can trigger hydra effects through the “prudent resource exploitation” mechanism. Furthermore, tension between mortality, foraging depression, and resource production predictably governs the range from strong cascades to hydra effects during epidemics.

## 2. Materials and methods

### (a) Experimental system

We parameterize and test our model in a planktonic system. We use isoclonal lines of the freshwater zooplankton host (*Daphnia dentifera*) to manipulate focal traits (Strauss *et al*. 2015). Hosts incidentally ingest free-living propagules of a virulent fungal parasite (*Metschnikowia bicuspidata*; Ebert 2005) while filter feeding on algal resources (*Ankistrodesmus falcatus*). Infected hosts have elevated death rate, and they release infectious propagules upon death. The short generation times of hosts, resources, and parasites enable multi-generation feedbacks during experimental epidemics.

### (b) Estimation of susceptibility and foraging traits

Susceptibility – the probability a host becomes infected given a single exposure – modulates population-level mortality from disease. For each of three clonal genotypes, we estimated susceptibility from the number of hosts becoming infected following controlled time and dose of propagules (Strauss *et al*. 2015). For these same host genotypes, we quantified depression of foraging rate when hosts contact spore propagules. Foraging rate of individual hosts was measured for eight hours at different spore concentrations (Strauss *et al*. 2019). Since terminal infection develops over longer time (~9 days, Stewart Merrill *et al*. 2019), the measured depression of foraging rate reflects behavior more than a symptom of infection. We fit foraging data for genotypes 1 and 2 (by Strauss *et al*. 2019, re-analyzed here) and for genotype 3 (unpublished) as an exponential function of spores *f*(*Z*) = *f*_0_e^−*αZ*^ (where *f*_0_ is the maximum foraging rate; see Appendix: section 2a). We determined significance of differences in these coefficients of foraging depression (*α*) by bootstrapping 95% CIs (Efron & Tibshirani 1993 see Appendix: section 2a for details). All analyses were performed in Rstudio (R Core Team 2019).

### (c) Resource-host-parasite model

We analyze a minimal model of logistically growing resources (*R*), susceptible hosts (*S*), infected hosts (*I*), and free-living parasite propagules (spores, *Z*). This model captures key biology of focal planktonic system (see also Table 1):

**Table 1.**
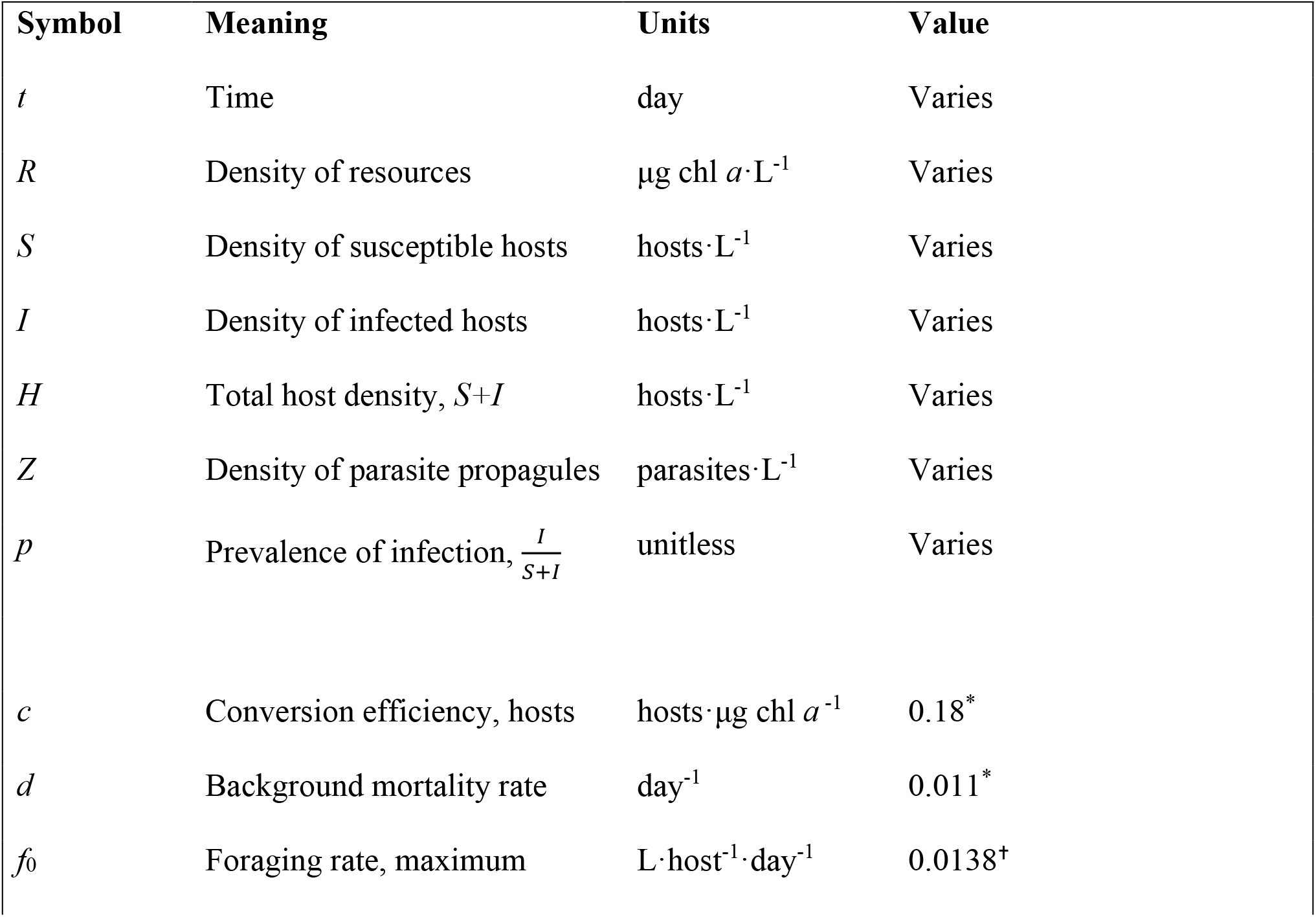
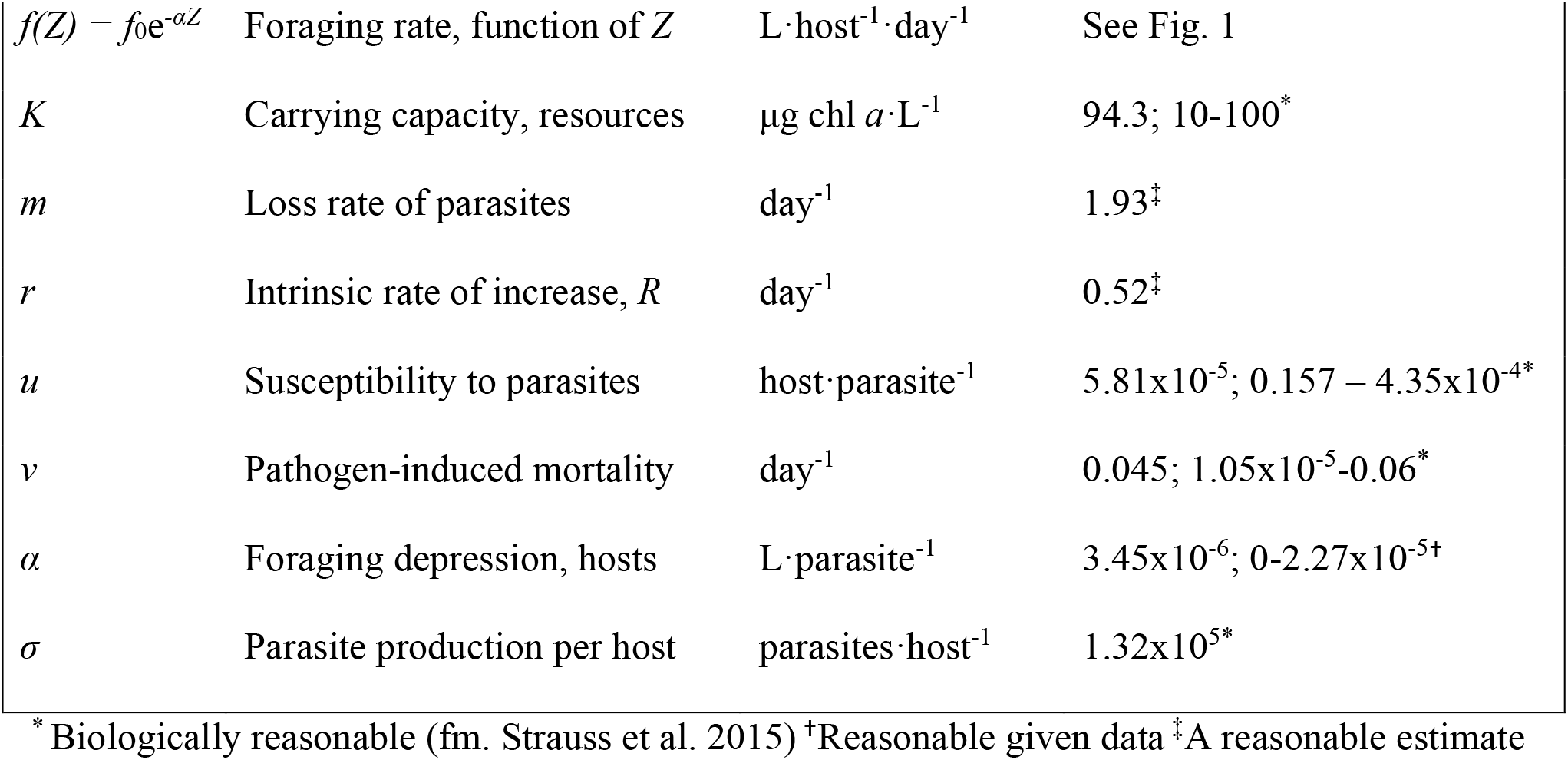
Symbols for state variables (top) and traits and other parameters (bottom) in the dynamical model (eq. 1). Default values are accompanied by ranges.

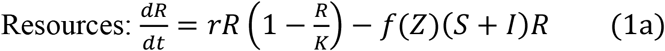

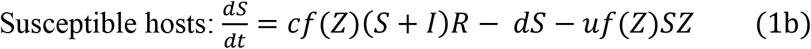

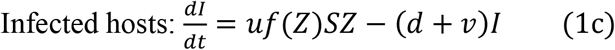

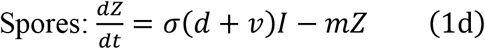

Resources grow logistically, a reasonable assumption allowing primary productivity to (potentially) increase during epidemics. Primary productivity is population-level recruitment of new resources per unit time: *rR*(1-*R*/*K*). Resource growth and production depend on intrinsic rate of increase *r* and carrying capacity *K* (first term, eq. 1a). Resources are consumed by susceptible (*S*) and infected (*I*) hosts foraging at rate *f*(*Z*) = *f*_0_*e*^*-αZ*^ (parameterized above) which declines exponentially with density of spores (*Z*); this exponential function prevents the unrealistic case of a negative feeding rate at high *Z*. This trait-mediated effect of parasites and may be strong (large *α*) or absent (*α* = 0). We assume equal foraging rate and foraging depression for susceptible and infected classes as a first, simplest approach (see Penczykowski *et al*. 2020 for a reduced foraging rate by infected hosts). Both host classes, *S* and *I*, convert resources into susceptible offspring (i.e., horizontal, environmental transmission; first term, eq. 1b; see Appendix: section 1b for reduced fecundity of infected hosts) with conversion efficiency *c*. For simplicity, infection does not lower fecundity. Total host density is *H* = *S* + *I*. Hosts experience background mortality at rate *d* (second term, eq. 1b) due to predation, senescence, and other factors. Susceptible hosts encounter parasites while foraging; their susceptibility, *u*, determines the proportion of hosts infected per encounter. Exposure and susceptibility jointly determine the transmission rate (often denoted *β*) from the environment to susceptible hosts, *uf*_0_*e*^−*αZ*^ (third term, eq. 1b). Infection converts susceptible hosts into infected hosts (first term, eq. 1c). These hosts suffer elevated death rate due to virulence of infection, *d*+*v* (where *v* is pathogen-induced mortality; second term eq. 1c). Death of infected hosts is the density-mediated effect of parasites. When infected hosts die, they release *σ* parasite propagules (*Z*) back into the environment (first term, eq. 1d). Losses of parasite propagules occur at background rate *m* (second term, eq. 1d), e.g., due to sinking, consumption (Strauss *et al*. 2015), solar radiation (Overholt *et al*. 2012), etc.

The model has a single, stable, endemic equilibrium for the range of parameter values considered. Since the exponential form of *f*_0_*e*^−*αZ*^ prevented general analytical solutions (though see eq. 2 for partial solutions), we found equilibrial densities at each parameter set with a root finder and evaluated stability of each using local stability analysis in Rstudio version 3.6.0 (R Core Team 2019). We then determine the effects of prevalence (*p*), susceptibility (*u*), foraging depression (*α*), and carrying capacity (*K*) on resource (*R*) and host (*H*) densities using a mixture of analytical and numerical techniques. We defined the strength of trophic cascades (resource release with host suppression) or hydra effects (resource release with host increase) as ratios of the stable endemic equilibrium to the disease-free boundary equilibrium. Since the mesocosm epidemics likely did not reach equilibria, we compared the closet analogue: we qualitatively matched them to transient dynamics of the model, and use the equilibria of the model to predict longer term outcomes. (See Appendix: section 1c for parameter values in which oscillations or bistability [when *α* > 0] arise).

### (d) Mesocosm test of model predictions

We test predictions of the dynamic model with populations of interacting algal resources, zooplankton hosts, and fungal parasites. To manipulate host traits, we stocked mesocosms with hosts of either one clonal genotype differing in *α* and *u* traits or a 50:50 mixture of two genotypes. In these mixtures, despite the potential for change in clone frequency, initial trait values qualitatively predicted patterns of density outcomes (see below); traits of mixed populations were bounded by those of the two genotypes and this is approximated by using the mean of the two genotypes’ traits. Using weighted average traits from genotype frequencies (not shown) did not improve model fit or qualitatively change any outcomes (using same methods as Walsman *et al*., in review). To manipulate carrying capacity of resources, we stocked algal resources into 50-liter mesocosms receiving either low or high nutrient supply (see Appendix: section 2b for more experimental details). Disease treatments were inoculated once with fungal spores (propagules). Altogether there were six susceptibility treatments (genotypes 1, 2, 3, 1&2, 1&3, or 2&3) × two nutrient supply treatments (low or high) × two parasite treatments (present or absent; each population treatment was replicated three times; 72 total mesocosms). With twice weekly sampling, we measured algal density, host density, and infection prevalence over c. seven host generations (~1 generation every 10 days; see Appendix: sections 2b, c).

We assume that temporal averages approximate equilibria from the model. In that model, connections between susceptibility (*u*) and prevalence (*p*) prove important mathematically. Hence, we first tested for their relationship (and with nutrient supply) using beta regression (see Appendix: section 2d). We next log_10_-transformed algal and host density, respectively, to disease, susceptibility, and nutrients. Log ratios reduce bias while having more normal error distributions (Hedges, Gurevitch & Curtis 1999). When disease drives a trophic cascade, it should have a significant positive effect on resource density and negative effect on host density. If susceptibility increases trophic cascade strength, there should be a significant positive interaction between susceptibility and disease for log_10_ resource density and negative interaction for log_10_ host density (see Appendix: section 2e). Additionally, we tested for a hydra effect in two genotype treatments at high nutrient supply using a nested ANOVA (see Appendix: section 2f). For each genotype, log_10_ host density was the response variable with mesocosms nested within disease treatment and repeated measurements nested within mesocosms. This nesting incorporated the power and autocorrelation of repeated measurements. Out of the original 72 mesocosms, nine were removed as outliers (most due to population extinction; see Appendix: section 2c). All analyses were performed in Rstudio (R Core Team 2019).

## 3. Results

### (b) Estimation of susceptibility and foraging traits

The range of trait values occupied by the three host genotypes facilitates manipulation of population-average traits. The three genotypes span almost an order of magnitude of susceptibility (Bristol 10: *u*_1_ = 5.81 × 10^−5^, A4-3: *u*_2_ = 1.09 × 10^−4^, Standard: *u*_3_ = 3.93 × 10^−4^). All host genotypes significantly depressed their foraging in the presence of parasites (decreasing curves in Fig. 1a; coefficients of foraging depression [*α*] with 95% CIs not overlapping zero; Fig. 1b). Foraging depression was more than twice as strong for genotype 1 (α = 3.5 × 10^−7^ L spore^−1^) as for genotypes 2 or 3 (1.4 × 10^−7^ and 7.6 × 10^−8^, respectively; 95% CIs of genotype 1 do not overlap those of the other two; Fig. 1b). Given this coefficient of foraging depression, the foraging rate of genotype 1 decreased by a factor of four over the range of spore doses in the trait assay (roughly corresponding to range of spore densities in the mesocosm, see Fig. S1).

**Figure 1.**
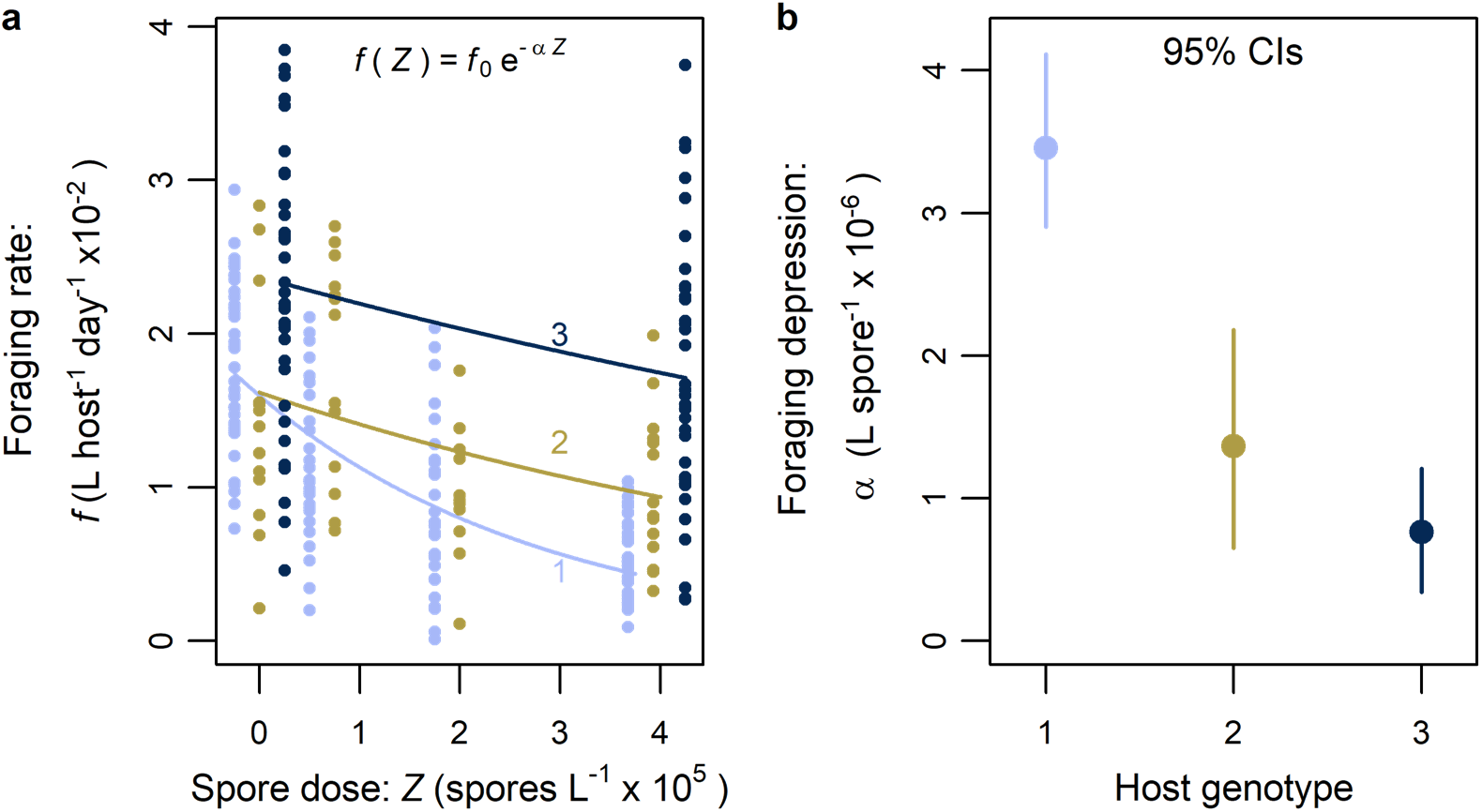
Differences among genotypes in the coefficient of foraging depression (α). (a) Fits of an exponential decay model for foraging rate with spores, *f*(*Z*) = *f*_0_ e^(-*αZ*)^ to data from a foraging experiment. Larger values of *α* indicate stronger depression of foraging rate with spore dose, *Z*. Points are jittered horizontally for clarity. (b) The coefficient of foraging depression (*α*) for genotype 1 was stronger than for other two genotypes (2 and 3). Error bars are boot-strapped 95% CIs.

### (c) Resource-host-parasite model

Susceptibility modulates host mortality while foraging depression (*α* > 0) enables the parasite-driven hydra effect. When host populations are more susceptible, they suffer higher prevalence of infection and thus higher mortality. When parasites drive higher mortality, they release resources and suppress host density more, producing a stronger trophic cascade. Hence, susceptibility acts analogously to attack rate of predators (see Appendix: section 1a and Shurin & Seabloom 2005). Yet, with lower susceptibility and mortality, foraging depression can drive a hydra effect. Insight arises from the numerator and denominator of host density (Schröder, van Leeuwen & Cameron 2014) derived from the resource equation (eq. 1a). Host density (*H*^*^ = *S*^*^+*I*^*^) is the ratio of primary productivity, *PP* = *rR*^*^(1-*R*^*^/*K*), to per host food consumption, *FC* = *f*(*Z*^*^)*R*^*^. Resource density, *R*^*^, in turn, is host death rate [*d* + *vp*^*^, with prevalence *p* = *I*^*\**^/*H*^*^] divided by fecundity per resource available [*cf*(*Z*^*^)]. Thus, host density depends on productivity and consumption of resources:

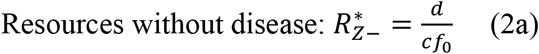

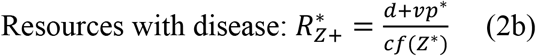

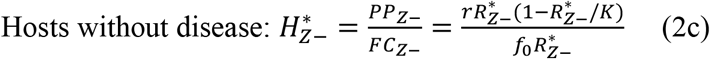

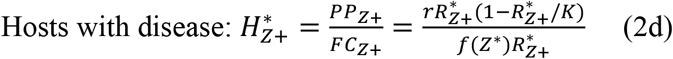

Because disease raises mortality, it still must increase the minimal resource requirement of hosts (compare eqs. 2a, b; *R*^*^_*Z+*_ *> R*^*^_*Z-*_since *f*(*Z*) < *f*_0_ and *d*+*vp*^*^ > *d*). Released resources increase or decrease primary productivity by drawing nearer to or farther from *K*/2 (because *PP is* maximized at *K*/2). *PP* drops when hosts had high minimal requirements without disease (*R*^*^_*Z-*_ > *K*/2). Lower *PP*, in turn, necessarily decreases host density (permitting only a cascade). But if hosts more strongly controlled resources without disease (*R*^*^_*Z-*_ < *K*/2), resource release can increase *PP* (if *R*^*\**^_*Z+*_ is closer to *K*/2 than *R*^*^_Z-_). However, increased *PP* alone may not suffice: hydra effects only emerge when increased *PP* (*PP*_*Z*+_ > *PP*_Z-_) outweighs increased food consumption per host (*FC*_*Z*+_ > *FC*_Z-_; eq. 2c, d).

Hence, food consumption places an additional constraint on the hydra effect. Food consumption per host (*FC*) is the product of foraging rate, *f*(*Z*^*^), and the density of resources, *R*^*\**^. Higher mortality from infection increases both *R*^*\**^ and *FC* when foraging rates stay constant [*α* = 0, so *f*(*Z*^*^) = *f*_0_]. If parasites only kill hosts (*v* > 0, *α* = 0), *FC* increases more with *R*^*^ than primary productivity can [since *H*^*^_*Z*+_ simplifies to *r*(1-*R*^*^/*K*) / *f*_0_, a decreasing function of *R*^*^]. So, parasites that only kill must decrease host density. If parasites only depress host foraging rate (*v* = 0, *α* > 0), resource density with disease simplifies to *R*^*^_*Z*+_ = *d*/[*cf*(*Z*^*^)] (from eq. 2b). This minimal resource requirement, *R*^*^_*Z*+,_ compensates for foraging depression completely; hence, *FC* remains constant (*FC*_*Z*+_ = *FC*_Z-_become equal when *v* = 0; eq. 2c, d). Thus, a parasite that only depresses foraging rate of hosts (but does not kill) will increase host density if primary productivity increases with disease. If parasites increase mortality and depress host foraging rate (*v* > 0, *α* > 0), they can drive a hydra effect if the increase in *PP* outweighs the simultaneous increase in *FC* per host. Thus, a tension between foraging depression and mortality emerges because they differentially influence production and consumption of resources.

These dueling effects of production and consumption explain why hydra effects become more likely with higher carrying capacity of the resource (*K*). Higher *K* elevates density of parasite propagules in the environment (*Z*) in both the mortality-only case (*v* > 0, *α* = 0; dashed red) and the foraging-depression case (*v* > 0, *α* > 0; solid red curve; Fig. 2a). More propagules, in turn, decrease foraging rate (when *α* > 0; Fig. 2b). Resources (*R*^*^) also increase with carrying capacity, but more so if hosts depress foraging (Fig. 2c). That larger release of resources can create a stronger increase in primary productivity - and more so at higher *K* (Fig. 2d). At the same time, food consumption per host (Fig. 2e) increases with *K* but less so with foraging depression than without. Hence, epidemics more easily drive a hydra effect at higher *K* (Fig. 2f).

**Figure 2.**
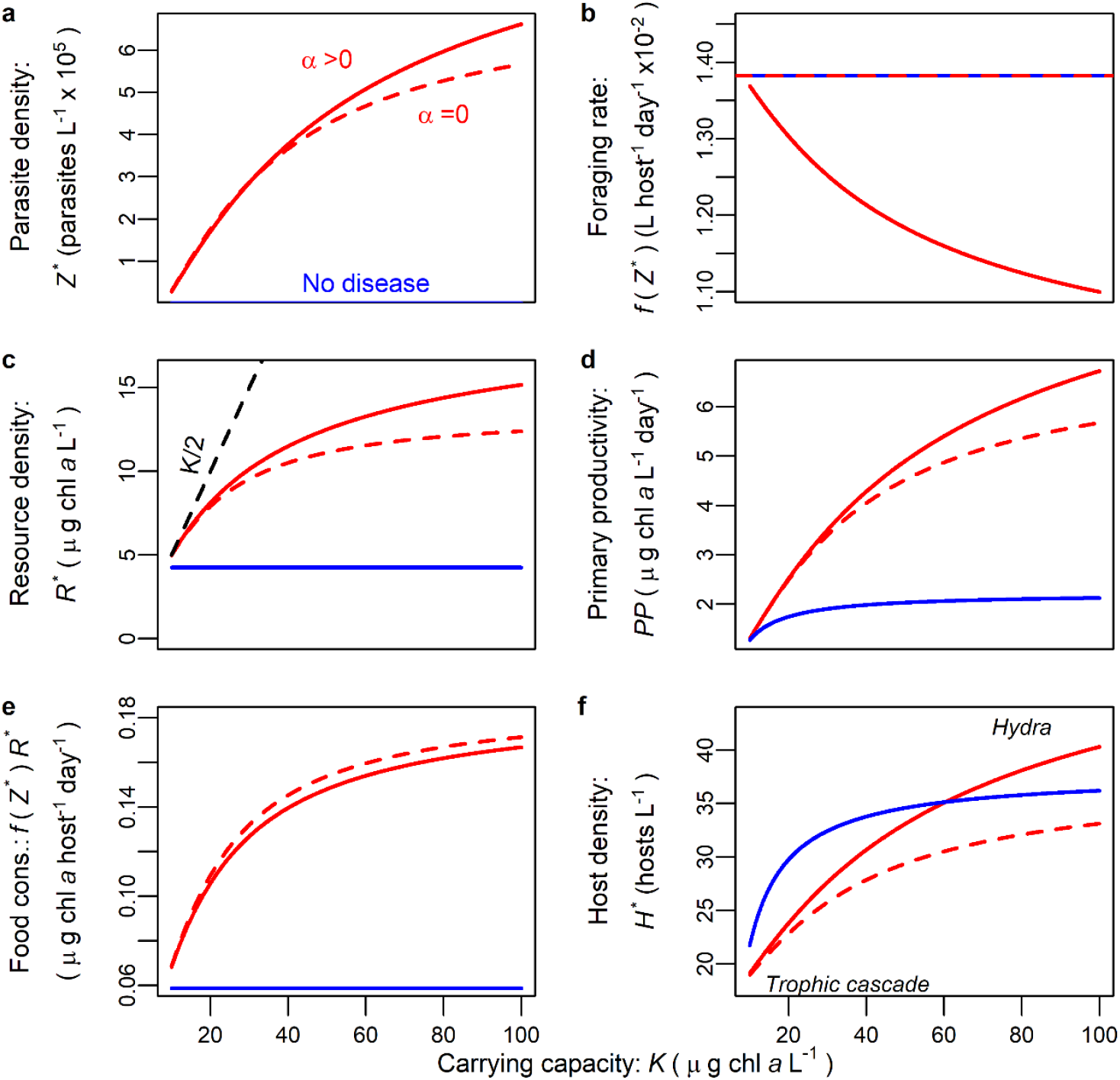
Foraging depression produces a hydra effect at high carrying capacity (see Fig. 3; eq. 2). Three cases shown: no-disease (blue, *Z*-), mortality-only (dashed red, *Z*+; *α* = 0, *v* > 0), and foraging depression (solid red, *Z*+; *α* > 0, *v* > 0). (a) Higher carrying capacity of the resource (*K*) increases propagule density (*Z*^*^), dropping (b) foraging rate if *α* > 0. (c) Parasites release resources (*R*^*^_*Z*+_ > *R*^*^_*Z*-_). *R*^*^_Z+_ is closer to *K*/2 (dashed black) with foraging depression. (d) Hence, primary productivity (*PP*) is highest when *α* > 0 and lowest without disease. (e) Food consumption, [*FC = f*(*Z*^*\**^)*R*^*\**^], is higher without foraging depression. (f) Given these responses of *PP* and *FC*, epidemics cause trophic cascades (*H*^*\**^_*Z*-_ > *H*^*\**^_*Z*+_ always) without foraging depression but can cause hydra effects when *α* > 0 at high *K*. (Parameters follow Table 1).

This framework predicts how carrying capacity (*K*), susceptibility (*u*), and foraging depression (*α*) jointly determine the range from strong cascades to hydra effects during epidemics. Although foraging rate drops with *α* and this releases resources, *K* has a stronger effect on resource release [indexed via ratio log_10_(*R*^*^_*Z+*_/*R*^*^_*Z-*_), eq. 2a, b; as parameterized, the contours are fairly vertical in Fig. 3a]. That resource release can then increase primary productivity. Foraging depression (high α) also decreases food consumption per host. Hence, the host ratio (*H*^*^_*Z+*_/*H*^*^_*Z-*_) increases with higher *K* (due to enhanced *PP*) and higher *α* (via lower FC), crossing from trophic cascade into hydra effect (past the black line [log_10_ ratio = 0] in Fig. 3b). That hydra effect becomes less likely when hosts experience higher mean per capita mortality. Mean mortality, *d* + *vp*, increases when susceptibility (higher *u*) raises prevalence (higher *p*). Higher *u* (hence higher mortality) releases resources strongly (larger resource ratios with higher *α* but especially higher *u*; Fig. 3c), boosting primary production. However, higher *u* also increases food consumption per host. Therefore, despite larger resource release, higher susceptibility makes hydra effects less likely (i.e., the region of host ratios sitting above the black line shrinks with higher *u*; Fig. 3d; see Fig. S2 for an exception at low *u* and high *α*). Similarly, if parasites exert higher pathogen-induced mortality, *v*, hydra effects become less likely (except at higher *α* and low *v*; see Fig. S3). Therefore, hydra effects arise at higher *K* and when hosts experience more foraging depression and less mortality. And trophic cascades are strongest when the combination of weak foraging depression and high susceptibility lead to high infection and mortality.

**Figure 3.**
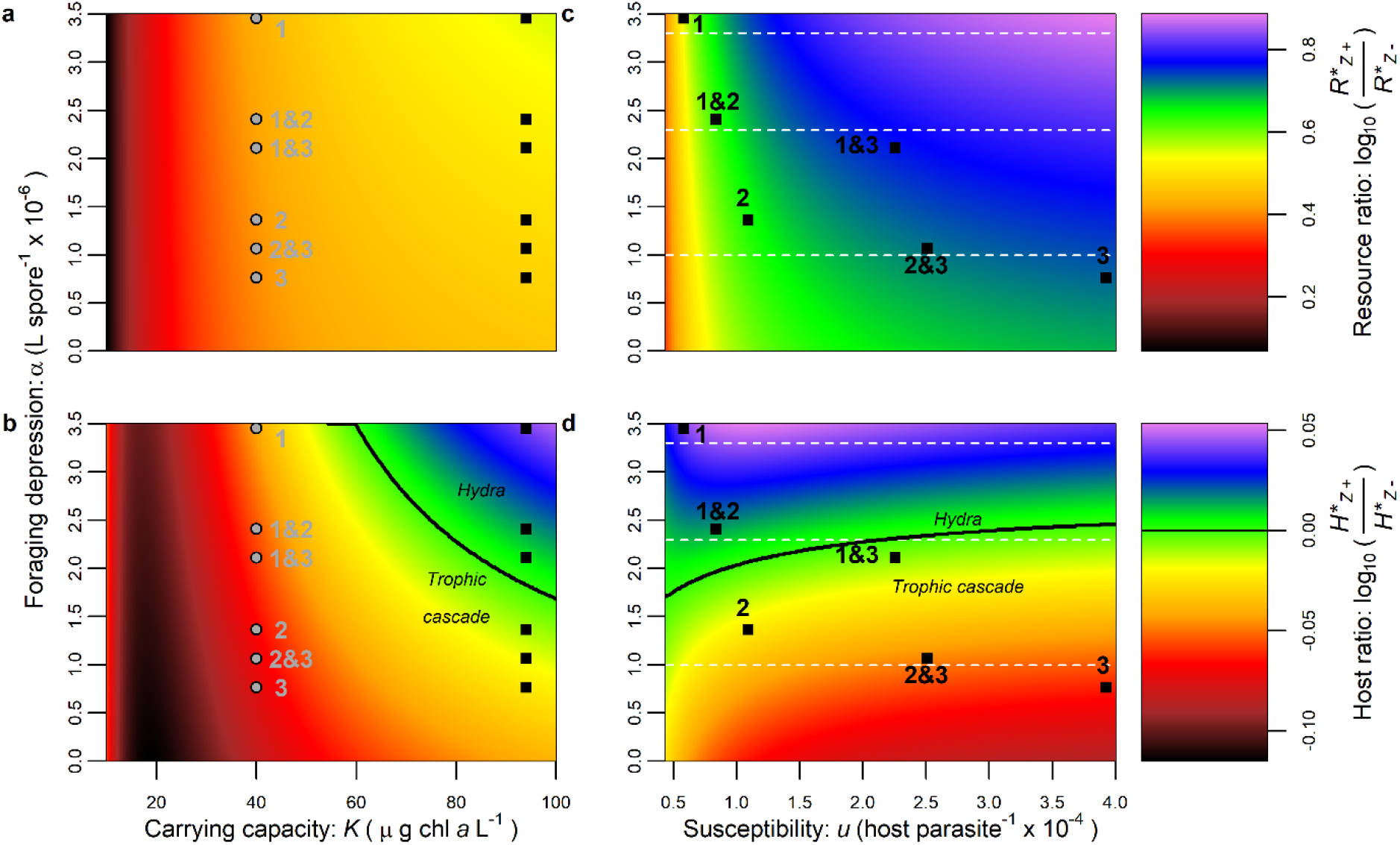
Combinations of carrying capacity (K, left column) or susceptibility (u, right) with foraging depression (α) create cascades or hydra effects. Points indicate traits of numbered genotype at low (light grey) or high (black) nutrient supply (note this comparison is qualitative). Colors indicate log_10_ density ratios of resource (top row: log_10_[*R*^*\**^_*Z*+_ /*R*^*\**^_*Z*-_]) or hosts (bottom: log_10_[*H*^*\**^_*Z*+_ /*H*^*\**^_*Z*-_]). Black curves note host ratio = 0 (above: hydra effects). *K-α space:* (intermediate *u* = 2.22 × 10^−4^) (a) Resource ratio increases with *α* but especially *K*. (b) Both *K* and *α* jointly increase host ratio, yielding a hydra effect at high enough *K*-*α* (see Fig. 2f). *u-α parameter space:* (high *K* = 94.3) Slices at dotted white lines shown in Fig. S2. (c) Higher *u* and α both increase resource ratio. (d) Higher susceptibility (*u*) increases parasite-driven mortality, reducing the region of hydra effects in *u*-*α* space. (Parameters follow Table 1).

These modeling results predict a qualitative pattern for the mesocosm results. The starting average traits (*u* and *α*) of mixed-genotype treatments are useful estimates for this qualitative prediction because the population is bounded within the traits of the two genotypes. More specifically, trait values (see Figure 3) indicate where genotype treatments should fall along the spectrum from strong cascades to hydra effects. These patterns calculated from equilibria are qualitatively similar when the system is not yet in equilibrium in simulations or mesocosms (see Figs 4, S4, S5, S6). Similarly, the presence of two host clones does not alter the predicted impact of host traits (Figs S4, S6). Therefore, the model predicts patterns linking traits to the cascade-hydra effect spectrum for the experiment to test.

**Figure 4.**
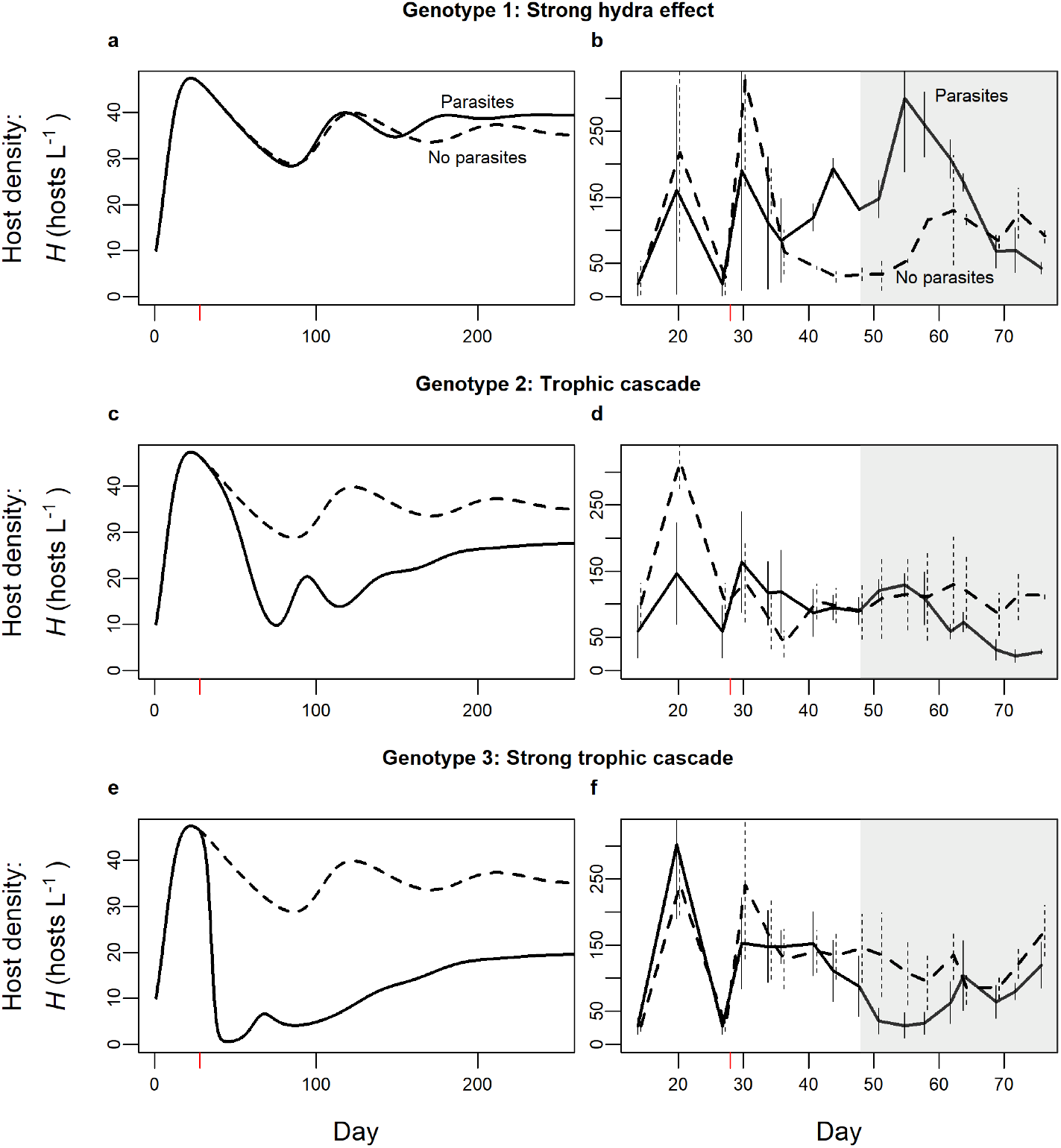
Simulated and experimental time series at high nutrients (K = 94.3 in simulations or 50 μg L^−1^ P in mesocosms) produce a spectrum ranging from hydra effects to trophic cascades. In both simulations and the experiment, hosts and parasites are added on days 0 and 28 (red tick mark), respectively. (a) With genotype 1’s traits, the hydra effect emerges given sufficient time as host density with parasites (solid) becomes higher than without (dashed). (b) Mesocosms of genotype 1 experienced a hydra effect [mean density across replicates with parasites (solid) or without (dashed), plotted at each time point; bars are standard error at each time point]. (c) With genotype 2’s traits, a trophic cascade (host suppression and resource release [Fig. S5]) occurs in simulations and (d) the mesocosm. (e-f) This cascade is larger for genotype 3 (the most susceptible). (Parameters follow Table 1). For analyses, average mesocosm density was taken from day 48 to 76 (gray region, see Methods for mesocosm). Experimental time series shifted slightly horizontally for clarity. Compare simulations to Fig. 3’s equilibrium outcomes and mesocosm time series to Fig. 5’s mesocosm averages.

### (d) Mesocosm test of model predictions

Most genotype treatments produced trophic cascades (released resources and suppressed host density). In populations with disease, higher susceptibility (*u*) increased prevalence of fungal infection (found from beta regression; P = 0.0067; see Fig. S1a and Appendix: section 2d). Higher nutrients also increased prevalence of disease (P = 0.0198), as predicted (see Fig. S1a). Relative to disease-free populations, disease released algal resources overall (P = 0.00126) and suppressed host density (P = 0.00137; see Table S1 for more model-data comparison). Susceptibility increased the strength of these trophic cascades, as predicted. Treatments with higher susceptibility displayed stronger release of algal resources (positive *u* × *Z+* interaction, P = 0.0277; see Figs S5d-f, S6d-f; increasing resource ratio in Fig. 5a). Susceptibility also strengthened host suppression (negative *u* × *Z*+ interaction, P < 0.001; Figs 4d-f, S4d-f; decreasing host ratio in Fig. 5b). These results match predictions for parasite-driven, density-mediated trophic cascades (see Figs 3c, d, 4a-c, S4a-c, S5a-c, S6a-c).

**Figure 5.**
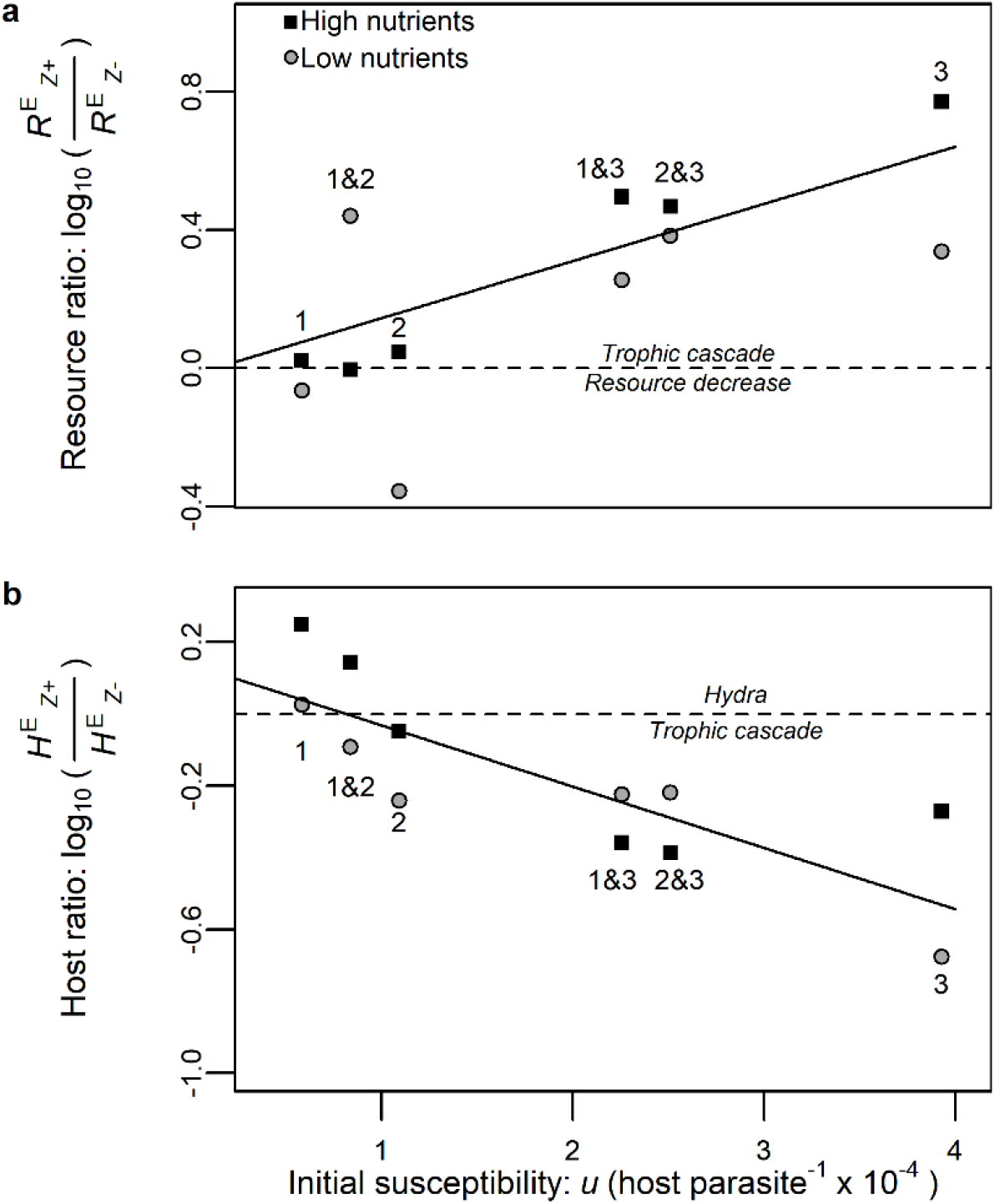
*In experimental epidemics, parasite drove cascades or hydra effects depending on susceptibility (u), foraging depression (α), and nutrient supply (K)*. Each point represents the average resource or host ratio. *Algal resources*: *R*^*E*^ *(E for ‘experiment’)*. (a) Higher susceptibility amplifies resource release (increasing log_10_ ratio of resources). (Error bars for ratios not shown: they are non-trivial to compute). *Plankton hosts: H*^*E*^. (b) Higher susceptibility amplifies trophic cascades (stronger host suppression; log_10_ host ratios become more negative with *u*). However, two genotype treatments with low susceptibility and strong foraging depression (1 and 1&2 together) displayed disease-driven hydra effects (log_10_ host ratios > 0) at high nutrient supply.

However, two sets of treatments displayed a hydra effect. At high nutrient supply, host density was higher with disease than without it for genotype 1 alone (strong hydra: 79% higher with disease, P = 0.00748; Figs 3d, 4a, and square labeled ‘1’ in 5b) and for the mixture of genotypes 1 & 2 (weaker hydra: 34% higher, P = 0.0201; Figs S4a and square ‘1&2’ in 5b). Since genotype 1 has low susceptibility (*u*) and high coefficient of foraging depression (α; Fig. 1), these results follow the model prediction for hydra effects at high *K*, lower *u*, and lower *α* (see Figs 3b,d, 4a, S4a).

## 4. Discussion

Our model qualitatively anticipated the strength of parasite-driven trophic cascades and hydra effects in a mesocosm experiment. Higher susceptibility of host genotypes increased prevalence of fungal infection, suppressed density of zooplankton hosts, and released algal resources. This result echoes how higher attack rate on prey strengthens predator-driven trophic cascades. However, at high nutrient supply, the least susceptible genotype experienced a parasite-driven hydra effect (higher host density during epidemics). This hydra effect arose partially because these treatments had lower susceptibility, hence lower per capita mortality rate. Vitally, these genotype treatments also showed strongest depression of foraging rate when encountering parasite propagules. Our model predicted that this combination of lower susceptibility and higher foraging depression could make parasites increase resource production (especially at high nutrients) more than food consumption during epidemics. Hence, this trait-environment combination predicted the strength of parasite-driven trophic cascades but also the appearance of hydra effects.

Susceptibility strengthened trophic cascades because it increased disease prevalence, hence parasite-driven mortality of hosts. We tested this prediction by manipulating susceptibility via a range of clonal genotypes of host populations. The susceptible genotypes, alone and in combination with each other, produced stronger parasite-driven trophic cascades (i.e., stronger host suppression and resource release). Parasite-driven trophic cascades are a well-established phenomenon (Buck & Ripple 2017); our model-experiment match adds this emerging area as the first to predict the strength of parasite-driven trophic cascades. It also demonstrates the power of the analogy to predator-driven cascades (Hall *et al*. 2008; Raffel, Martin & Rohr 2008 and see Appendix: section 1a for more details). Like in the disease case, higher mortality inflicted by predators should lead to stronger cascades. Hence, predator-driven cascades should be stronger when predators capture prey more quickly (analogous to higher transmission rate and related higher susceptibility; Shurin & Seabloom 2005). Additionally, factors that enhance attack rate – e.g., predators’ body size (Jochum *et al*. 2012; Simonis 2013), metabolic rate (Borer *et al*. 2005), and feeding behavior (Katano, Natsumeda & Suguro 2013) - each increase cascade strength via mortality effects (Vucic-Pestic *et al*. 2011; Katano, Natsumeda & Suguro 2013; Simonis 2013). Thus, both susceptibility to infection and attack rate of predators increase mortality of victims. Therefore, both drive stronger cascades of their victims.

However, the combination of model and experiment revealed another possibility: epidemics of virulent parasites can also produce hydra effects. The hydra emerges because disease can alter two components of host density, production and per capita consumption of resources (Schröder, van Leeuwen & Cameron 2014). Both production and consumption, in turn, depend on mortality, foraging depression, and the minimum resource requirement (*R*^*^) of hosts. To start, virulent effects of infection lead to higher host *R*^*^ (as parasites often do; Holdo *et al*. 2009; Buck & Ripple 2017). When hosts strongly control their resources (i.e., low enough *R*^*^ without disease), increased *R*^*^ can elevate resource production (Fourqurean *et al*. 2010). All else equal, this higher production could support higher host density. However, increased mortality increases per-capita food consumption of hosts. If parasites only kill hosts but do not change foraging rate, food consumption always increases more than resource production (in our model). These parasites can only cause trophic cascades (see Buck & Ripple 2017 for many examples). In contrast, a hydra effect can arise when parasites depress foraging rate of hosts. Foraging depression, a trait-mediated effect, arises commonly in this host-parasite system (Hite *et al*. 2017; Strauss *et al*. 2019) and others (Hite, Pfenning & Cressler 2020) and in predator-prey systems (Morgan 1988; Laundré, Hernández & Ripple 2010). Here, foraging depression enables production to increase more than per capita consumption of resources during epidemics. In such circumstances, foraging depression can create hydra effects.

Both the dynamical model and the experiment anticipate conditions needed for a parasite-driven hydra effect. Hosts with low susceptibility (low *u*) suffer less infection and infection-induced mortality (pathogen-induced mortality). Lower mortality coupled with stronger depression of foraging rate (higher *α*) can cause resource release without huge increases in food consumption. Furthermore, resource release can elevate primary productivity more at higher carrying capacity (when *R*^*^ of the host without disease is less than *K*/2). The experiment verified these predictions; hydra effects arose for hosts having both low susceptibility and strong foraging depression in environments with high supply of nutrients (high *K*). This example with parasites adds to a few other examples of hydra effects caused by mortality fixed by the experiment and falls under a “prudent resource exploitation” mechanism (Abrams 2009; Schröder, van Leeuwen & Cameron 2014). Previously, others have shown how parasites can increase biomass (Ohlberger *et al*. 2011; Preston & Sauer 2020) or both body mass and survival for a specific life stage of hosts (Washburn & Mercer 1991). Total host density also increased in large fungal epidemics in this host in lake populations (Penczykowski *et al*. 2020). Here, in a controlled experiment, parasites increased total host density. That outcome emerged due to dynamical feedbacks from interacting hosts, resources, and parasites over multiple generations.

Given its emergence here, how can we anticipate parasite-driven hydra effects in other systems? What factors might constrain or amplify their possibility? First, the relationship between susceptibility (*u*) and foraging depression (*α*) traits may make hydra effects more or less likely. In the set of genotypes here, susceptibility and foraging depression were negatively correlated, yielding strong trophic cascades (high *u*, low *α*; lower right Fig. 3d) or strong hydra effects (low *u*, high *α*; upper left Fig. 3d). For another set of genotypes of this host, these traits did not correlate negatively (Strauss *et al*. 2019). A positive correlation might reduce the probability of strong cascades or strong hydra effects. Second, hosts may evolve weaker or stronger foraging depression as selection weighs lowered fecundity against lowered exposure to parasites. Third, hydra effects should be somewhat less likely when parasites impose more harm to host fitness. For systems where foraging depression does not reduce parasite exposure, exposure and parasite-induced mortality would remain high, making a hydra effect somewhat less likely. Similarly, parasites imposing high virulence on mortality and on fecundity are less likely to drive hydra effects. Castrators, for example, should more likely cause cascades (see Appendix: section 1b). Future work could clarify the importance of each of these factors for hydra effects in this and other systems.

Here, using models and experiments, we delineate when parasites cause trophic cascades or hydra effects. Parasite-driven trophic cascades have emerged in various systems. Here, we develop and test a trait-based model framework experimentally. Yet, the parasite-mediated hydra effect that arose reveals an even newer possibility: parasites that kill hosts could increase host density if they depress foraging. This finding seems baffling until one embraces feedbacks between hosts and resources. Now disease ecologists can ask: are disease-mediated hydra effects rare? Or could they arise more commonly - if we just know where and when to look for them? Here, the mathematical model guides us. First, parasites must not increase host mortality too greatly. Second, hosts must depress their foraging rate in the presence of parasites (a trait-mediated effect). Third, higher resource density must increase primary productivity. If resources cannot respond dynamically to host density or if productivity does not increase with higher resources, hydra effects cannot occur via this prudent resource exploitation mechanism. Nonetheless, we show how these three factors (mortality, foraging depression, productivity) govern the range from strong parasite-driven trophic cascades to hydra effects.

## Acknowledgements

O. Schmidt and J. Obergfell provided assistance with the experiments. C. Lively, F. Bashey-Visser, M. Wade, A. Ramesh, and T. Deblieux provided valuable feedback on the manuscript. This work was supported by NSF DEB 1353749, 1655656, and an NSF GRFP 1342962 to J. Walsman.

## Author contributions

JW and SH designed the mesocosm experiment, AS and SH designed the trait measurements, AS performed the trait measurements, JW performed the mesocosm experiment, mathematical modeling, data analysis, and wrote the first draft of the manuscript while all authors contributed substantially to revisions.

## References

Abrams, P.A. (2009) When does greater mortality increase population size? The long history and diverse mechanisms underlying the hydra effect. Ecology Letters, 12, 462–474.

Beckerman, A.P., Uriarte, M. & Schmitz, O.J. (1997) Experimental evidence for a behavior-mediated trophic cascade in a terrestrial food chain. Proceedings of the National Academy of Sciences, 94, 10735–10738.

Boivin, G., Hance, T. & Brodeur, J. (2012) Aphid parasitoids in biological control. Canadian Journal of Plant Science, 92, 1–12.

Borer, E.T., Seabloom, E.W., Shurin, J.B., Anderson, K.E., Blanchette, C.A., Broitman, B., Cooper, S.D. & Halpern, B.S. (2005) What determines the strength of a trophic cascade? Ecology, 86, 528–537.

Buck, J., Weinstein, S. & Young, H. (2018) Ecological and evolutionary consequences of parasite avoidance. Trends in ecology & evolution, 33, 619–632.

Buck, J.C. & Ripple, W.J. (2017) Infectious agents trigger trophic cascades. Trends in ecology & evolution, 32, 681–694.

Cleaveland, S., Laurenson, M. & Taylor, L. (2001) Diseases of humans and their domestic mammals: pathogen characteristics, host range and the risk of emergence. Philosophical Transactions of the Royal Society of London. Series B: Biological Sciences, 356, 991–999.

Cooper, J., Crawford, R.J., De Villiers, M.S., Dyer, B.M., Hofmeyr, G.G. & Jonker, A. (2009) Disease outbreaks among penguins at sub-Antarctic Marion Island: a conservation concern. Marine Ornithology, 37, 193–196.

Daszak, P., Cunningham, A.A. & Hyatt, A.D. (2000) Emerging infectious diseases of wildlife--threats to biodiversity and human health. science, 287, 443–449.

Dobson, A., Lafferty, K.D., Kuris, A.M., Hechinger, R.F. & Jetz, W. (2008) Homage to Linnaeus: how many parasites? How many hosts? Proceedings of the National Academy of Sciences, 105, 11482–11489.

Dwyer, G. & Elkinton, J.S. (1993) Using simple models to predict virus epizootics in gypsy moth populations. Journal of Animal Ecology, 1–11.

Ebert, D. (2005) Ecology, epidemiology, and evolution of parasitism in Daphnia. National Library of Medicine (US), National Center for Biotechnology Information, Bethesda, MD, United States.

Ebert, D., Lipsitch, M. & Mangin, K.L. (2000) The effect of parasites on host population density and extinction: Experimental epidemiology with Daphnia and six microparasites. American Naturalist, 156, 459–477.

Efron, B. & Tibshirani, R.J. (1993) An Introduction to the Bootstrap. Chapman & Hall, New York.

Fourqurean, J.W., Manuel, S., Coates, K.A., Kenworthy, W.J. & Smith, S.R. (2010) Effects of excluding sea turtle herbivores from a seagrass bed: overgrazing may have led to loss of seagrass meadows in Bermuda. Marine Ecology Progress Series, 419, 223–232.

Fry, W.E. & Goodwin, S.B. (1997) Resurgence of the Irish potato famine fungus. BioScience, 47, 363–371.

Hall, S.R., Becker, C.R., Duffy, M.A. & Cáceres, C.E. (2011) Epidemic size determines population-level effects of fungal parasites on Daphnia hosts. Oecologia, 166, 833–842.

Hall, S.R., Lafferty, K.D., Brown, J.H., Cáceres, C.E., Chase, J.M., Dobson, A.P., Holt, R.D., Jones, C.G., Randolph, S.E. & Rohani, P. (2008) Is infectious disease just another type of predator-prey interaction. Infectious Disease Ecology: the effects of ecosystems on disease and of disease on ecosystems, 223–241.

Hedges, L.V., Gurevitch, J. & Curtis, P.S. (1999) The meta-analysis of response ratios in experimental ecology. Ecology, 80, 1150–1156.

Hite, J.L., Penczykowski, R.M., Shocket, M.S., Griebel, K.A., Strauss, A.T., Duffy, M.A., Cáceres, C.E. & Hall, S.R. (2017) Allocation, not male resistance, increases male frequency during epidemics: a case study in facultatively sexual hosts. Ecology, 98, 2773–2783.

Hite, J.L., Pfenning, A.C. & Cressler, C.E. (2020) Starving the enemy? Feeding behavior shapes host-parasite interactions. Trends in ecology & evolution, 35, 68–80.

Hochachka, W.M. & Dhondt, A.A. (2000) Density-dependent decline of host abundance resulting from a new infectious disease. Proceedings of the National Academy of Sciences, 97, 5303–5306.

Holdo, R.M., Sinclair, A.R., Dobson, A.P., Metzger, K.L., Bolker, B.M., Ritchie, M.E. & Holt, R.D. (2009) A disease-mediated trophic cascade in the Serengeti and its implications for ecosystem C. PLoS Biol, 7, e1000210.

Horan, R.D. & Fenichel, E.P. (2007) Economics and ecology of managing emerging infectious animal diseases. American journal of agricultural economics, 89, 1232–1238.

Jochum, M., Schneider, F.D., Crowe, T.P., Brose, U. & O’Gorman, E.J. (2012) Climate-induced changes in bottom-up and top-down processes independently alter a marine ecosystem. Philosophical Transactions of the Royal Society B: Biological Sciences, 367, 2962–2970.

Katano, O., Natsumeda, T. & Suguro, N. (2013) Diurnal bottom feeding of predator fish strengthens trophic cascades to benthic algae in experimental flow-through pools. Ecological Research, 28, 907–918.

Laundré, J.W., Hernández, L. & Ripple, W.J. (2010) The landscape of fear: ecological implications of being afraid. The Open Ecology Journal, 3.

Matsuda, H. & Abrams, P.A. (2004) Effects of predator prey interactions and adaptive change on sustainable yield. Canadian Journal of Fisheries and Aquatic Sciences, 61, 175–184.

Morgan, M.J. (1988) The influence of hunger, shoal size and predator presence on foraging in bluntnose minnows. Animal Behaviour, 36, 1317–1322.

Ohlberger, J., Langangen, Ø., Edeline, E., Claessen, D., Winfield, I.J., Stenseth, N.C. & Vøllestad, L.A. (2011) Stage-specific biomass overcompensation by juveniles in response to increased adult mortality in a wild fish population. Ecology, 92, 2175–2182.

Overholt, E.P., Hall, S.R., Williamson, C.E., Meikle, C.K., Duffy, M.A. & Caceres, C.E. (2012) Solar radiation decreases parasitism in Daphnia. Ecology Letters, 15, 47–54.

Penczykowski, R.M., Hall, S.R., Shocket, M.S., Ochs, J.H., Lemanski, B.C., Sundar, H. & Duffy, M.A. (2020) Virulent disease epidemics can increase host density by depressing foraging of hosts. bioRxiv.

Philpott, S.M., Maldonado, J., Vandermeer, J. & Perfecto, I. (2004) Taking trophic cascades up a level: behaviorally-modified effects of phorid flies on ants and ant prey in coffee agroecosystems. Oikos, 105, 141–147.

Preston, D.L. & Sauer, E.L. (2020) Infection pathology and competition mediate host biomass overcompensation from disease. Ecology, 101.

R Core Team (2019) R: A language and environment for statistical computing. R Foundation for Statistical Computing, Vienna, Austria.

Raffel, T.R., Martin, L.B. & Rohr, J.R. (2008) Parasites as predators: unifying natural enemy ecology. Trends in ecology & evolution, 23, 610–618.

Raveh, A., Kotler, B.P., Abramsky, Z. & Krasnov, B.R. (2011) Driven to distraction: detecting the hidden costs of flea parasitism through foraging behaviour in gerbils. Ecology Letters, 14, 47–51.

Roelke-Parker, M.E., Munson, L., Packer, C., Kock, R., Cleaveland, S., Carpenter, M., O’brien, S.J., Pospischil, A., Hofmann-Lehmann, R. & Lutz, H. (1996) A canine distemper virus epidemic in Serengeti lions (Panthera leo). Nature, 379, 441.

Schröder, A., van Leeuwen, A. & Cameron, T.C. (2014) When less is more: positive population-level effects of mortality. Trends in ecology & evolution, 29, 614–624.

Shurin, J.B. & Seabloom, E.W. (2005) The strength of trophic cascades across ecosystems: predictions from allometry and energetics. Journal of Animal Ecology, 74, 1029–1038.

Simonis, J.L. (2013) Predator ontogeny determines trophic cascade strength in freshwater rock pools. Ecosphere, 4, 1–25.

Stewart Merrill, T.E., Hall, S.R., Merrill, L. & Cáceres, C.E. (2019) Variation in immune defense shapes disease outcomes in laboratory and wild Daphnia. Integrative and Comparative Biology, 59, 1203–1219.

Strauss, A.T., Bowling, A.M., Duffy, M.A., Cáceres, C.E. & Hall, S.R. (2018) Linking host traits, interactions with competitors and disease: mechanistic foundations for disease dilution. Functional Ecology, 32, 1271–1279.

Strauss, A.T., Civitello, D.J., Cáceres, C.E. & Hall, S.R. (2015) Success, failure and ambiguity of the dilution effect among competitors. Ecology Letters, 18, 916–926.

Strauss, A.T., Hite, J.L., Civitello, D.J., Shocket, M.S., Cáceres, C.E. & Hall, S.R. (2019) Genotypic variation in parasite avoidance behaviour and other mechanistic, nonlinear components of transmission. Proceedings of the Royal Society B, 286, 20192164.

Thrall, P. & Burdon, J. (2000) Effect of resistance variation in a natural plant host–pathogen metapopulation on disease dynamics. Plant Pathology, 49, 767–773.

Vredenburg, V.T., Knapp, R.A., Tunstall, T.S. & Briggs, C.J. (2010) Dynamics of an emerging disease drive large-scale amphibian population extinctions. Proceedings of the National Academy of Sciences, 107, 9689–9694.

Vucic-Pestic, O., Ehnes, R.B., Rall, B.C. & Brose, U. (2011) Warming up the system: higher predator feeding rates but lower energetic efficiencies. Global Change Biology, 17, 1301–1310.

Washburn, J.O. & Mercer, D.R. (1991) Regulatory role of parasites: impact on host population shifts with resource availability. science, 253, 185–188.

Wood, C.L., Byers, J.E., Cottingham, K.L., Altman, I., Donahue, M.J. & Blakeslee, A.M. (2007) Parasites alter community structure. Proceedings of the National Academy of Sciences, 104, 9335–9339.

