## Supporting Information for "Parasite-driven cascades or hydra effects: susceptibility and foraging depression shape parasite-host-resource interactions"

### Section 1: Additional model details

#### *(a) Mortality-only model of predator-driven, density-mediated trophic cascades*

Mortality-only parasite-driven cascades (eq. 1 with $\alpha$ = 0; Fig. S1) follow patterns nearly identical to those of predator-driven cascades. A model of a logistically growing resource ($R$), a prey ($S$) that consumes the resource, and a predator ($P$) that consumes the prey provides a tractable comparison:

$$\frac{dR}{dt} = rR\left(1 - \frac{R}{K}\right) - f_0 SR \qquad \text{(S1a)}$$

$$\frac{dS}{dt} = cf_0SR - dS - f_PSP \qquad \text{(S1b)}$$

$$\frac{dP}{dt} = c_Pf_PSP - d_PP \qquad \text{(S1c)}$$

As in the model for parasites, resources grow logistically with intrinsic rate of increase $r$ and carrying capacity $K$ (first term, eq. S1a). Resources are consumed by prey ($S$, to mirror the model); these prey forage at per-capita rate $f_0$ (second term, eq. S1a). Consumed resources are converted into prey with efficiency $c$ (first term, eq. S1b). Prey die at background per-capita rate $d$ (second term, eq. S1b), and predators eat them at per-capita attack (capture) rate, $f_P$ (third term, eq. S1b). Consumed prey are converted into predators with efficiency $c_P$ (first term, eq. S1c). Predators die at background per-capita rate $d_P$. This minimal model assumes linear functional forms for clear analytical and dynamical interpretation – and for easier comparison to the disease analogue here. Other predator-prey-resource models consider further biological detail, such as Type II functional responses and metabolic types (Shurin & Seabloom 2005).

Due to their structural similarities, this Predator model yields predictions analogous to those of parasite-driven trophic cascades. Carrying capacity ($K$) has the same interpretation as a driver of resource productivity in both models (since it pertains to a logistically growing resource in both cases); susceptibility ($u$) acts similarly to attack rate of predators ($f_P$). Therefore, it is useful to determine the effects of carrying capacity ($K$) and attack rate of predators on resource and prey density without ($R^*_{P-}$, $S^*_{\mathrm{P-}}$) and with ($R^*_{P+}$, $R^*_{P+}$, respectively) predators, and on the ratios of resources ($R^*_{P+}$/ $R^*_{P-}$) and prey ($S^*_{P+}$/ $S_{P-}$) with and without predators:

$$R^*_{P-} = \frac{d}{cf_0} \qquad \text{(S2a)}$$

$$R^*_{P+} = K(1 - \frac{f_o d_P}{c_P f_P r}) = \frac{d + f_P P^*}{cf_0} \qquad \text{(S2b)}$$

$$\frac{R^*_{P+}}{R^*_{P-}} = \frac{cf_0K}{d}\left(1 - \frac{f_0 d_p}{c_p f_p r}\right) = \frac{d + f_P P^*}{d} \qquad \text{(S2c)}$$

$$S^*_{P-} = \frac{r}{f_0}\left(1 - \frac{d}{cf_0K}\right) = \frac{r}{f_0K}(K - R^*_{P-}) \qquad \text{(S2d)}$$

$$S^*_{P+} = \frac{d_P}{c_P f_P} = \frac{r}{f_0}\left(1 - \frac{d + f_P P^*}{cf_0K}\right) = \frac{r}{f_0K}(K - R^*_{P+}) \qquad \text{(S2e)}$$

$$\frac{S^*_{P+}}{S^*_{P-}} = \frac{cf_0^2 K d_p}{c_p f_p r(cf_0K - d)} = \frac{K - R^*_{P+}}{K - R^*_{P-}} \qquad \text{(S2f)}$$

$$P^*_{P+} = \frac{cf_0K\left(1 - \frac{f_0 d_p}{c_P f_P r}\right) - d}{f_P} = \frac{cf_0}{f_P}(R^*_{P+} - R^*_{P-}) \qquad \text{(S2g)}$$

Predators ($P$) increase prey mortality, thus they increase the minimal resource requirements ($R^*$) of their prey ($S$). Therefore, resource density with predators ($R^*_{P+}$) must be higher than without predators ($R^*_{P-}$;,compare eq. S2a and S2b). Carrying capacity ($K$) does not increase resources without predators because $R^*_{P-}$ is the ratio of background mortality ($d$) to per-resource fecundity ($cf_0$; eq. S2a). With predators, $R^*_{P+}$ is linearly proportional to $K$. Biologically, $R^*_{P+}$ increases with predators ($P^*$; eq. S2g) because higher mortality increases minimum resource requirement of the prey (eq. S2b). Thus, $K$ amplifies how much predators release resources ($d/dK\ [R^*_{P+}/R^*_{P-}] > 0$; eq. S2c). Additionally, attack rate of the predator ($f_P$) does not affect resources without predators (eq. S2a) but increases it with them (eq. S2b). Thus, $f_P$ amplifies how much predators release resources ($d/df_P\ (R^*_{P+}/R^*_{P-}) > 0$; eq. S2c). Overall, resource release by predators with $K$ and $f_P$ in this model and a more complex one (Shurin & Seabloom 2005) mirror the (indirect) effects of parasites on resources of hosts.

Because they increase minimal resource requirements of their prey, predators suppress density of their prey. Prey density with predators ($S^*_{P+}$, proportional to $K - R^*_{P+}$) must be lower than that without predators ($S^*_{P-}$, proportional to $K - R^*_{P-}$; compare eqs. S2d and S2e). Carrying capacity ($K$) increases prey without predators because prey enjoy top-down control of resources (eq. S2d). With predators, prey become fixed at the minimum prey requirement of predators and therefore cannot increase with $K$ (eq. S2e). Thus, $K$ amplifies how much predators harm prey density (d/d$K$ [$S^*_{P+}/S^*_{P-}$] < 0 always; eq. S2f). Increasing attack rate of predators ($f_P$) decreases prey density with predation (since $f_p$ increases $R^*_{P+}$; eq. S2e). Thus $f_P$ amplifies how much predators suppress prey (d/d$f_P$ [$S^*_{P+}/S^*_{P-}$] < 0; eq. S2f). Consequently, the effects of predators on prey density via attack rate $f_P$ mirror those of parasites on host density via susceptibility $u$ in the mortality-only model case. Such analogous predictions only hold for carrying capacity when considering resource release. The victim suppression response instead differs between predators and prey (see below).

***(b) Virulence on fecundity (θ) and cascades vs. hydra effects***

Virulence on fecundity should accentuate trophic cascades. Many parasites reduce host fecundity (for several examples, see Ebert, Lipsitch & Mangin 2000), ranging from partial to full castration (no fecundity from infected hosts). The model can be easily altered to incorporate the possibility of fecundity reduction:

$$\frac{dR}{dt} = rR\left(1 - \frac{R}{K}\right) - f(Z)(S + I)R \qquad \text{(S4a)}$$

$$\frac{dS}{dt} = cf(Z)(S + \theta I)R - dS - uf(Z)SZ \qquad \text{(S4b)}$$

$$\frac{dI}{dt} = uf(Z)SZ - (d + v)I \qquad (S4c)$$

$$\frac{dZ}{dt} = \sigma(d + v)I - mZ \qquad \text{(S4d)}$$

The update introduces $\theta$, relative fecundity of infected hosts. If $\theta = 1$, infected hosts have full fecundity (i.e., the assumption made before). If $\theta = 0$, parasites castrate fully: infected hosts consume resources [at rate $f(Z)$] but produce no offspring (hence, this model is not quite like the predator-prey analogue; eq. S1). Fecundity reduction ($\theta$) then alters equilibrium densities or resources, $R^*_{Z+}$, and hosts, $H^*_{Z+}$ during epidemics (from eqs. 2b, d to S5):

$$R^*_{Z+} = \frac{d + vp}{cf(Z^*)(1 - p + \theta p)} = \frac{d + vp}{cf(Z^*)(1 - p) + \theta cf(Z^*)p} \qquad \text{(S5a)}$$

$$H^*_{Z+} = \frac{r}{f(Z^*)}\left(1 - \frac{R^*_{Z+}}{K}\right) \qquad \text{(S5b)}$$

where minimal resource requirement of hosts during epidemics, $R^*_{Z+}$, is the ratio of mortality ($d + vp$) to per resource fecundity (eq. S5a). However, notice that per resource fecundity now differs for susceptible hosts, $cf(Z)$, and infected hosts, $cf(Z)\theta$, (weighted by frequencies 1-$p$ and $p$). Host density, $H^*_{Z+}$ (eq. S5b) remains similar before (compare to eq. 2d), but with an updated value of $R^*_{Z+}$ (eq. S5a; compare denominator to denominator in eq. 2b). We do not fully analyze this model but provide intuitive predictions to be tested by future studies.

Fecundity reduction seems likely to reduce the possibility of a hydra effect compared to the virulence on mortality. Fecundity reduction (represented by $\theta < 1$) reduces host fitness overall, likely despite a small compensatory decrease in infection prevalence ($dp^*/d\theta < 0$; if hosts evolve, the compensatory decrease in prevalence is stronger). Reduced host fitness increases the resources required for the host population to survive during epidemics (increases $R^*_{Z+}$, compared to eq. 2b). Increased $R^*_{Z+}$, all else equal, must increase food consumption more than it increases primary productivity (because food consumption increases linearly with $R^*$ while $PP$ increases less than linearly, the reasoning why mortality alone must suppress host density). Thus, stronger castration (decreasing $\theta$) should further suppress host density (as found theoretically and empirically by Ebert, Lipsitch & Mangin 2000 but without a dynamic resource). However, decreasing $\theta$ could make a hydra effect more likely if, through an added mechanism (not considered in eq. S4), parasite density ($Z^*$) increases and depresses host foraging rate enough. Such a mechanism could involve gigantism: parasites that strongly or fully castrate (lower $\theta$) increase parasite production (higher $\sigma$) by directing host energy toward parasites (as in Ebert *et al.* 2004; Hall, Becker & Cáceres 2007).

Overall, it seems unlikely that virulence on fecundity would completely undermine the pattern of hydra effects to trophic cascades across genotypes. Strong foraging depression and low susceptibility populations suffer low prevalence of infection so virulence on fecundity will have a small impact, leaving hydra effects largely intact. Weak foraging depression and high susceptibility populations suffer high prevalence so virulence on fecundity should amplify resource release and host suppression, strengthening cascades. Virulence on fecundity may decrease prevalence, making these patterns somewhat more complex; such feedback on prevalence would have to be surprisingly strong to completely undermine the pattern across genotypes. Detailed modeling of these feedbacks is beyond the scope of this paper.

***(c) Outcomes other than one stable equilibrium in the main model (eq. 1)***

*Oscillations:* The model analyzed in the main text can produce oscillations instead of a stable interior equilibrium whether or not foraging depression occurs. These oscillations can arise when the carrying capacity of the resource ($K$) is very high (e.g., all parameters default except $K$ = 377). Weakening the negative density dependence of the resource, which is usually a stabilizing factor, is likely involved but other feedback loops may also be important. Foraging depression tends to reduce the possibility for oscillations, but they may still arise (e.g., high $K$ = 472 and weak $\alpha = 3.455 \times 10^{-7}$, all others as default, see Table 1).

*Multiple stable equilibria:* All plots presented in the main text use parameter ranges that give only one stable equilibrium. In contrast, foraging depression ($\alpha > 0$) does allow two simultaneously stable equilibria. Positive feedbacks between parasites and host density can create alternative stable states. For some parameter values (e.g. $c = 22.5$, $d = 0.0176$, $f_0 = 0.0161$, $K = 265$, $m = 9.59$, $u = 1.66 \times 10^{-4}$, $v = 0.19$, $w = 8.39$, $\alpha = 7.50 \times 10^{-8}$, $\sigma = 7.75 \times 10^{4}$), two endemic equilibria can be stable. A low disease equilibrium has lower parasite density, higher foraging rate, lower resource density, lower primary productivity, and lower host density. If instead parasites are denser, host foraging rate is strongly depressed (because $\alpha > 0$), resources are denser and more productive and thus support a larger host population with lower prevalence. These alternative stable states arise because of positive feedbacks between parasite density and host density. If hosts have low density, parasites will be sparse, and hosts will have a high foraging rate. This high foraging rate keeps host density low by overgrazing resources. If hosts become denser, however, parasite density increases, depressing host foraging rate and increasing host density. Further theoretical exploration could more clearly demonstrate the feedbacks, biological feasibility, and dynamical implications of this bistability.

***(d) Mortality-only case of the disease model: comparison to predator-driven cascades – Fig. S1, Table S1***

To study the morality-only case of the disease model, we needed to use numerical approaches. Intuitively, higher susceptibility ($u$) and carrying capacity ($K$) should lead to higher prevalence of infection ($p$). Also, it seems that higher $K$ should increase host density with disease ($H^*_{Z+}$; as long as $K$ doesn't increase $p^*$ very fast: see eq. S3e). Because the expressions involved are very large, we evaluated equilibrium quantities along broad parameter ranges, from $10^{-2}$ to

$10^2$ x default parameter values (see Table 1). We divided these ranges into $10^4$ evenly spaced values, then used a Latin Hypercube search (McKay, Beckman & Conover 2000) to find equilibrium densities at each parameter combination. At each parameter set, we increased or decreased $K$ or $u$ 10% to determine its effect on equilibria, without ($\alpha = 0$) and with foraging depression ($\alpha > 0$). Without foraging depression, higher $K$ or $u$ always increased prevalence, and higher $K$ always increased host density with disease. With foraging depression, prevalence can decrease with $K$ (but only increased in our focal parameter range: see Figures and Table 1). Additionally in seven of $10^4$ parameter sets, $u$ decreases prevalence (e.g., $c = 1.318302 \times 10^{1}$, $f_0 = 9.865147 \times 10^{-1}$, $\alpha = 3.061821 \times 10^{-04}$, $u = 3.486 \times 10^{-06}$, $d = 3.702714 \times 10^{-1}$, $v = 3.087158$, $s = 3.247983 \times 10^{6}$, $r = 1.916043 \times 10^{1}$, $K = 9.410805 \times 10^{3}$, $m = 1.191349 \times 10^{2}$). Additionally, in 1 of $10^4$ parameter sets, $K$ decreases host density with disease (e.g., $c = 8.051979$, $f_0 = 1.338437$, $\alpha = 2.285137 \times 10^{-04}$, $u = 4.82811 \times 10^{-06}$, $d = 2.285037 \times 10^{-1}$, $v = 2.215772 \times 10^{-1}$, $s = 3.624676 \times 10^{6}$, $r = 1.253682 \times 10^{-1}$, $K = 8.735303 \times 10^{3}$, $m = 1.290146 \times 10^{2}$). So, in the vast majority of cases and always within the biologically relevant range of parameter values, the intuitive effects of $K$ and $u$ on prevalence and host density hold.

With their relationships to prevalence established, we then evaluated the effects of $u$, $K$, and parasites on resource and host density with or without disease. With those densities, we calculated ratios (and related $\log_{10}$ of density ratios), common metrics of cascade strength (Shurin *et al.* 2002; Shurin & Seabloom 2005). With these metrics, cascades become stronger with smaller (log) ratio of hosts and higher (log) ratio of resources, i.e., with stronger host suppression and resource release, respectively. For the simple case where $\alpha = 0$ (see eq. 2 more generally), these quantities are:

$$R^*_{Z-} = \frac{d}{cf_0} \qquad (S3a)$$

$$R^*_{Z+} = \frac{d + vp^*}{cf_0} \qquad \text{(S3b)}$$

$$\frac{R^*_{Z+}}{R^*_{Z-}} = \frac{d + vp^*}{d} \qquad \text{(S3c)}$$

$$H^*_{Z-} = \frac{r}{f_0}\left(1 - \frac{d}{cf_0K}\right) = \frac{r}{f_0}\left(1 - \frac{R^*_{Z-}}{K}\right) \qquad \text{(S3d)}$$

$$H^*_{Z+} = \frac{r}{f_0}\left(1 - \frac{d + vp^*}{cf_0K}\right) = \frac{r}{f_0}\left(1 - \frac{R^*_{Z+}}{K}\right) \qquad \text{(S3e)}$$

$$\frac{H^*_{Z+}}{H^*_{Z-}} = \frac{cf_0K - d - vp^*}{cf_0K - d} \qquad \text{(S3f)}$$

Because parasites increase host mortality, they increase the minimum resource density required by hosts (a ratio of losses to per resource gains of hosts) from $R^*_{Z-}$ (eq. S3a) to $R^*_{Z+}$ (eq. S3b). Carrying capacity ($K$) does not increase resource density without disease (eq. S3a; superimposed red and blue dashed lines; Fig. S1c). However, with disease, $K$ increases prevalence ($p^*$) and thus $R^*_{Z+}$ (eq. S3b; red solid curve above blue Fig. S1c). Susceptibility ($u$) does not affect resource density without disease (dashed lines are flat; Fig. S1c) but increases resource density with disease by elevating prevalence (solid curves increase with $u$; Fig. S1c). Hence, the resource density ratio increases with both carrying capacity and susceptibility [eq. S3c; $\mathrm{d}/\mathrm{d}K\,(R^*_{Z+}/R^*_{Z-}) > 0$, red curve sits above the blue one; $\mathrm{d}/\mathrm{d}u\; R^*_{Z+}/R^*_{Z-} > 0$, curves increasing with $u$-axis: Fig. S1d]. Stated simply, both higher $u$ and $K$ lead to larger resource release.

In this mortality-only model case, parasites can only suppress host density. Mathematically, mortality increases the minimum resource requirement, $R^*_{Z+} > R^*_{Z-}$, so host density declines $H^*_{Z+} < H^*_{Z-}$ (see eqs. S3d, e; solid line below dashed one in Fig. S1e). Carrying capacity ($K$) increases host density without or with parasites (both $\mathrm{d}H^*_{Z-}/\mathrm{d}K > 0$ [analytically] and $dH^*_{Z+}/dK > 0$ [in numerical searches]; dashed red curve lies above dashed blue [Fig. S1e]). However, $K$ may increase or decrease the host ratio (the increase example where $K$ weakens host suppression, $d/dK\ [H^*_{Z+}/H^*_{Z-}] > 0$, is shown with red curve above blue in Fig. S1f). Susceptibility ($u$) does not affect host density without disease (eq. S3d; flat $H^*_{Z-}$ curves, Fig. S1e), but decreases it with disease through increasing prevalence ($p^*$; eq. S3e; decreasing $H^*_{Z+}$ curves, Fig. S1e). So, higher susceptibility increases host suppression (eq. S3f decreases; curves decreasing in Fig. S1f). Summarizing, in the mortality-only model case, parasites suppress host density more strongly when hosts have higher susceptibility but not necessarily when carrying capacity is higher. Allowing foraging depression in the parasite model ($\alpha > 0$) does not qualitatively change these patterns for hosts and resources.

The mortality-only model case shows that parasite-driven, density-mediated trophic cascades should function mostly like predator-driven, density-mediated ones. Parasites that only increase mortality can only suppress host density and release resources. The resource release (higher resource ratio, $R^*_{Z+}/R^*_{Z-}$) and host suppression (lower host ratio, $H^*_{Z+}/H^*_{Z-}$) both become stronger with higher susceptibility, $u$ (analogous to predator attack rate). Resource ratio also increases with carrying capacity, $K$, in both disease and predator-driven cascades. However, host ratio increases or decreases with $K$; its analogue only decreases with $K$ in predator-driven cascades. This difference arises because infected hosts accumulate with $K$ whereas predators fix prey density at their minimal requirement, a value unchanging with $K$. Predators fix density of their prey at their minimal prey requirement: $S^*_{P+} = d_P/[c_P f_P]$; eq. S2e). In contrast, host density ($S$+$I$) still increases with $K$ during epidemics due to accumulation of infected hosts ($I$), even though the parasite itself has a minimal requirement for susceptible hosts itself ($S^*_{Z+} = m / [u f_0 \sigma]$, a ratio of losses to gains like that of the predator). Because this accumulation of $I$ allows host density ($S$+$I$) to increase with $K$, host ratio (eq. S3f) can increase or decrease with $K$. The case with predators is simpler: captured prey are immediately removed by predators, so prey ratio (eq. S2f) can only decrease with $K$. Otherwise, the mortality-only case (eq. 1 with $\alpha = 0$) and predator model (eq. S1) produce analogous predictions for strength of trophic cascades. Thus, when parasites only kill, density-mediated trophic cascades largely resemble those with predators, yielding predictable effects of susceptibility, nutrient supply, and disease in the experiment (Table S1).

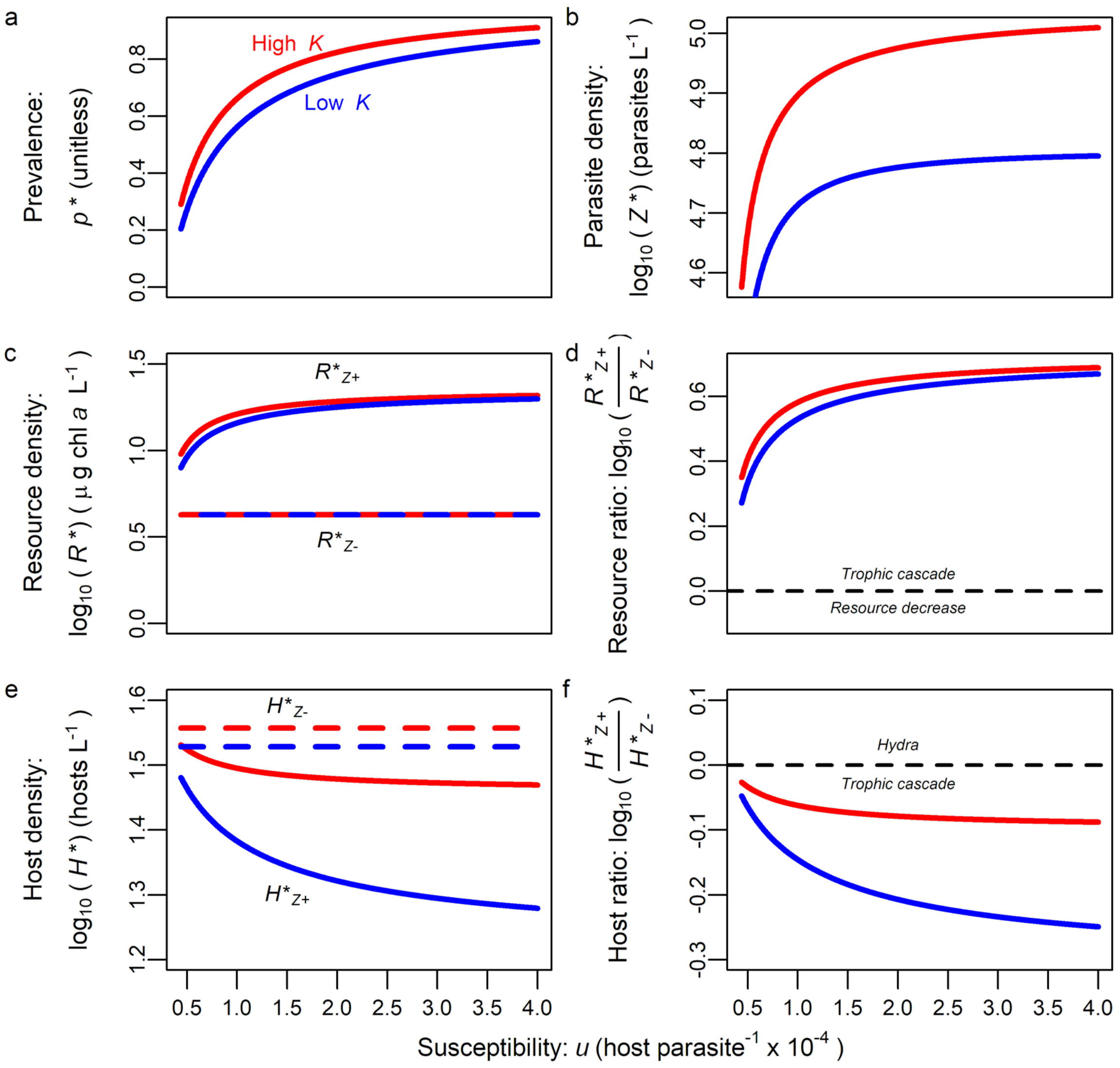

**Figure S1.** *Predicted effects of susceptibility on cascade strength at equilibrium in mortality-only case (eqs. 1, S3).* (a) Higher susceptibility ($u$) and carrying capacity ($K$) both lead to higher prevalence ($p^*$) of infection (b) and increased parasite density ($Z^*$). (c) Without disease (dashed lines), resource density is fixed at the minimum requirement of the host ($R^*_{Z-}$) and unaffected by susceptibility, $u$, or carrying capacity ($K$). With disease (solid curves), resources ($R^*_{Z+}$) increase above $R^*_{Z-}$ with $u$ and $K$. (d) Resource ratio: densities of resources with and without disease. Values above zero on a $\log_{10}$ scale indicate trophic cascade; below zero, resources decrease (which cannot occur in this model). Resource ratio increases with both $u$ and $K$ (both curves increasing and red curve above blue). (e) Without disease, susceptibility does not affect host density ($H^*_{Z-}$). With disease (solid), total host density ($H^*_{Z+}$) decreases with susceptibility. Higher $K$ leads to higher host densities (red curves above corresponding blue). (f) Host ratio: densities of hosts with and without disease ($H^*_{Z+} / H^*_{Z-}$). Values below zero on a log scale indicate trophic cascade; above zero indicate a hydra effect. Host ratio decreases with $u$ (both curves decreasing) but not necessarily with $K$ (e.g., blue curve below red in this example). Together, (a), (d), and (f) show that trophic cascade strength increases with $u$ as disease spreads more easily. $K$ = 20 (low) or 94.3 (high) µg chl $a$/L; other parameter values listed in Table 1.

***(e) Hydra effects and cascades with susceptibility (u) – Fig. S2 showing slices of α in Fig. 4c, d***

Increasing susceptibility to infection can promote disease, counterintuitively increasing host density if it amplifies a hydra effect. Generally, increasing $u$ amplifies the negative impact of parasites on host populations (host suppression). But, if there is already a hydra effect, increasing $u$ can amplify that hydra effect. Increasing susceptibility, $u$, increases parasite density, $Z^*$, particularly with higher levels of foraging depression, $\alpha$ (Fig. S2a). Therefore, foraging rate of hosts drops with $u$ (more $Z$) and $\alpha$ (stronger sensitivity to $Z$; Fig. S2b). Since epidemics become larger with $u$, resource density (i.e., the host's minimal requirement) increases with $u$. It also increases further with sensitivity of foraging depression (Fig. S2c). If the minimal resource requirement of the host without disease ($R^*_{Z-}$) lies below $K/2$ (density at peak resource productivity), the increase in resource density with disease (to $R^*_{Z+}$) can increase productivity of the resource (where again, $PP = r R^* (1 - R^*/K)$; Figs S2d, 2). Parasites that depress host foraging, then, have a stronger effect on $PP$ than food consumption. Notice, however, that the effects of foraging depression on per host food consumption, $f(Z^*)$ $R^*_{Z+}$ almost completely cancels; food consumption increases more with $u$ than with $\alpha$ (Fig. S2e). Host density ($H^*$; Fig. S2f) is the ratio of *PP* to food consumption (Fig. 2). Two patterns emerge. First, hydra effects are more likely at higher $\alpha$ because it increases *PP* with small effect on consumption. Conversely, when $\alpha$ is small, the *PP* boost from epidemics is smaller, too small to cause a hydra effect given the increase in consumption (so a cascade arises instead). At intermediate $\alpha$, we find a shift with increasing $u$ from hydra effect to cascade (as consumption increases faster with $u$ than *PP*). If hosts do not depress their foraging rate ($\alpha = 0$, or if $R^*_{Z-} > K/2$), then increasing susceptibility always decreases host density. Second, when a hydra effect is possible, it may be strongest (i.e., peak in host ratio) at intermediate $u$. At this level of $u$, parasite propagules ($Z$) become dense enough to reduce foraging rate while not adding too much mortality for hosts.

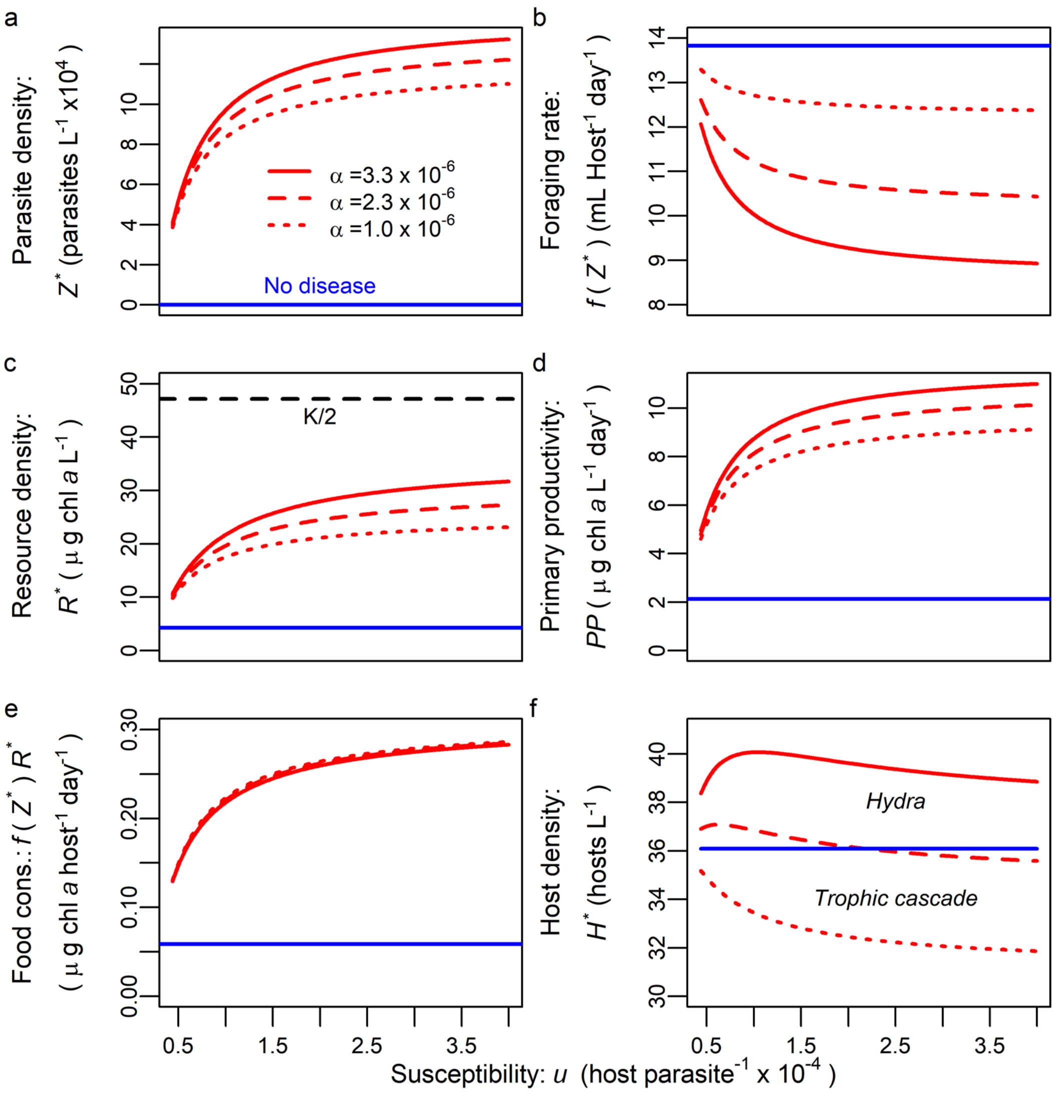

**Figure S2.** *Hydra effects and cascades with susceptibility (u) and different foraging depression (α).* Values of foraging depression (contours) here correspond to horizontal slices of Figs 4c, d. (a) Higher susceptibility ($u$) leads to higher parasite density with disease (red) but has no effect without it (blue). Higher foraging depression ($\alpha$) can increase parasite density ($Z^*$) due to associations with host density (red contours). (b) Higher $Z$ with increasing susceptibility depresses foraging rate, $f(R^*)$, particularly when $\alpha$ is larger. (c) Resource density, $R^*$, increases with susceptibility as more hosts are infected (higher $p^*$ with $u$) and each host forages less.

Increased resources ($R^*_{Z+}$) can be closer to $K/2$ (dashed black line). (d) Because $R^*_{Z+}$ becomes closer to $K/2$, primary productivity, *PP*, increases with $u$ and $\alpha$. (e) Food consumption, $f(Z^*)R^*$, rises with higher $u$ to compensate for more mortality but increases less for higher $\alpha$. (f) If foraging depression is strong enough (solid and dashed red), higher susceptibility can lead to increased host density ($H^*_{Z+} > H^*_{Z-}$, hence a hydra). Even so, host density reaches a maximum at intermediate susceptibility. Higher susceptibility decreases foraging rate slightly (via increased $Z^*$; panel b) but increases mortality. Thus, increased susceptibility can drive a transition from hydra effect to trophic cascade (dashed red). [$\alpha = 3.3 \times 10^{-6}$ (solid), $2.3 \times 10^{-6}$ (dashed), $1.0 \times 10^{-6}$ (dotted); see Table 1 for other parameter values].

#### *(f) Higher virulence (v) and cascades vs. hydra effects – Fig. S3*

The outcome of trophic cascade or hydra effect depends on a tension between mortality and foraging depression. Higher virulence mortality ($v$) of parasites increases direct harm to host fitness, more strongly increasing resource density (i.e., the minimal resource requirement of hosts, $R^*_{Z+}$ [eq. 2b]; Fig. S3a). Higher foraging depression also increases resource density (Fig. S3a; hence, resource ratio increases up and to the right). Higher virulence tends to depress host density (eq. 2d; Fig. S3b; host ratio declines to the right). Increasing foraging depression increases host density and can drive a hydra effect (Fig. S3b; trophic cascade below black curve and hydra effect above). With higher virulence, stronger foraging depression is required to still give a hydra effect (black line increasing in Fig. S3b). Once in $v$-$\alpha$ space producing a hydra effect, a different pattern can arise. At high $\alpha$, increasing virulence can sometimes amplify an existing hydra effect (host ratio increasing with $v$ for $\alpha = 3.5 \times 10^{-6}$). Here, higher virulence increases conversion of infected hosts into parasites, which depress foraging rate. (This result assumes that propagule yield per infected host, $\sigma$, would not change with $v$, an unlikely assumption biologically). With high enough $\alpha$ but not too high (not shown), this extra foraging depression can increase host density further. Generally, however, higher mortality virulence decreases host density because $v$ increases food consumption more than primary productivity. Furthermore, in a numerical search of equilibrium densities, higher $v$ never increases host density in the mortality-only model case ($v$> 0 but $\alpha$ = 0).

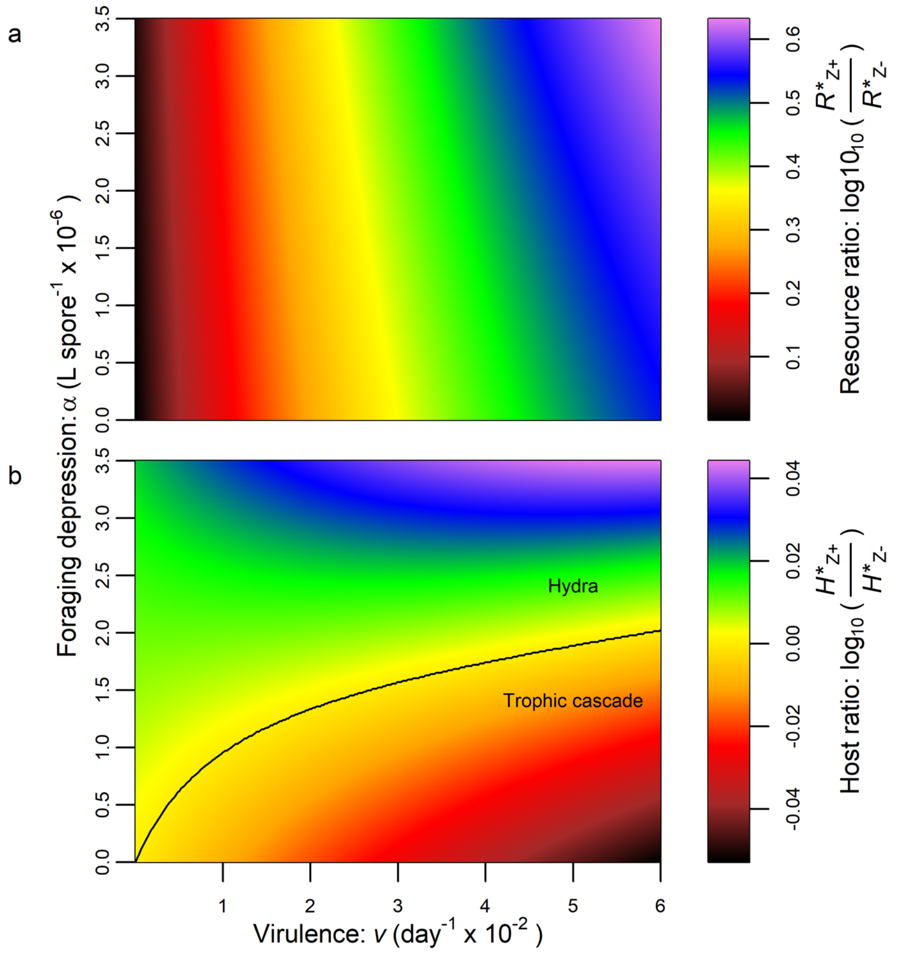

**Figure S3.** *Higher virulence (added mortality from infection) makes it harder for parasites to drive a hydra effect.* Mortality virulence ($v$ > 0) kills more hosts, leading to a trophic cascade, while foraging depression can increase productivity enough to overwhelm this mortality effect and drive a hydra effect. (a) Both virulence and foraging depression increase resource release ($\log_{10}$ of resource ratio, $R^*_{Z+}$ / $R^*_{Z-}$). (b) Higher virulence, $v$, generally suppresses host density (decreases $\log_{10}$ of host ratio $H^*_{Z+}$ / $H^*_{Z-}$) while foraging depression increases host ratio. Hydra effects (above black line), therefore, are more likely for less virulent parasites that depress foraging [black line, where $\log_{10}(H^*_{Z+} / H^*_{-}) = 0$, increases with $v$]. See Table 1 for parameter values.

***(g) Time series of model simulations and mesocosm data***

We also run simulations with two genotypes to match two-genotype populations. We do this simply with a host population composed of a 50:50 ratio of individuals with the traits of the two clones (see Discussion for future exploration of evolution of these traits). Thus, average foraging rate for a population is given by $f_{av} = f_0[\exp(-\alpha_1 Z) + \exp(-\alpha_2 Z)]/2$ while average transmission rate is given by $\beta_{av} = f_0(u_1 e^{-\alpha 1Z} + u_2 e^{-\alpha 2Z})/2$. With populations of two clones, simulations still adhere closely to the equilibrium, model patterns (see Figs S4, S6).

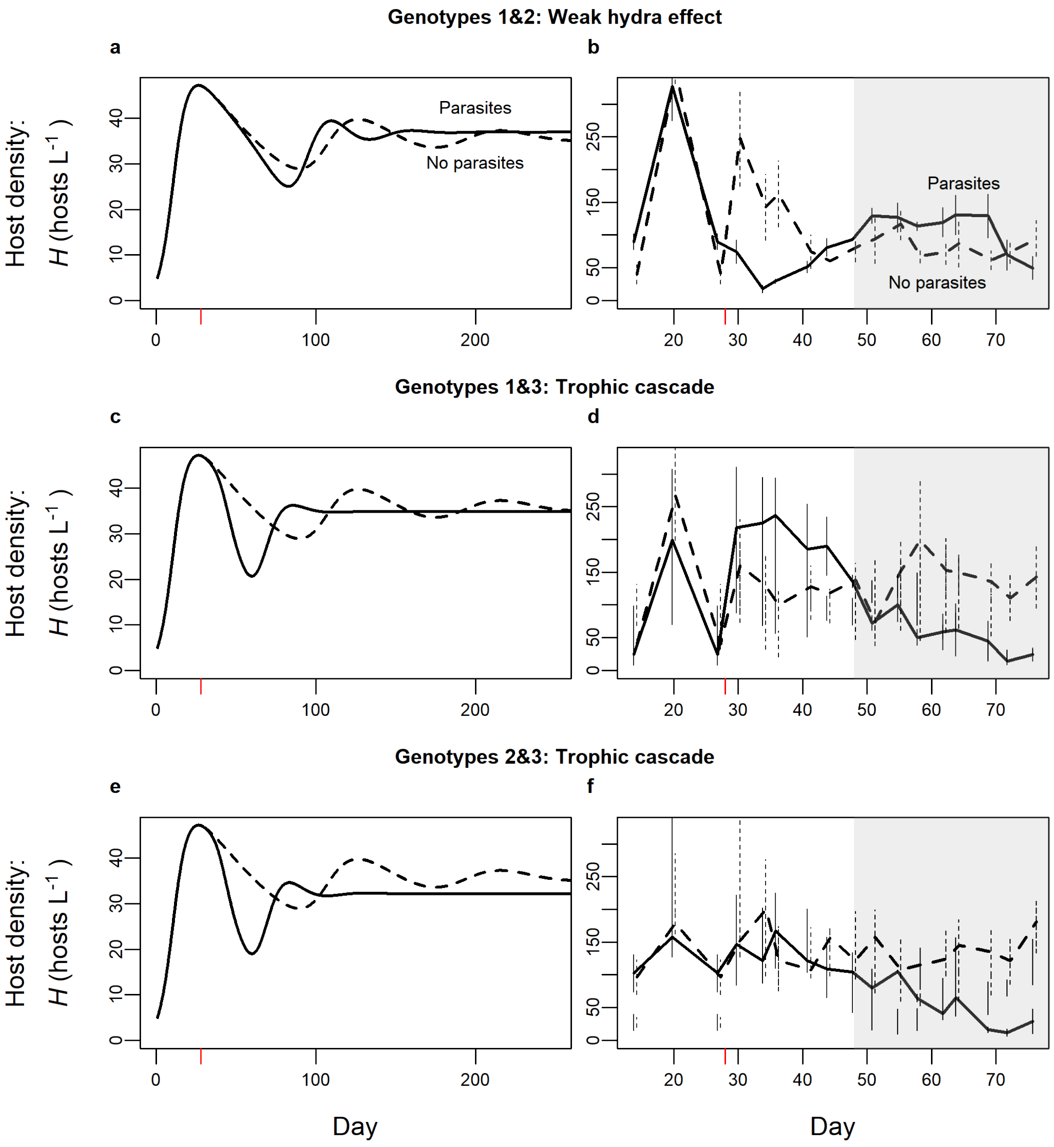

330

331 **Figure S4.** *Simulated and experimental time series at high nutrients (*K *= 94.3 in simulations or*

332 *50 µg L*$^{-1}$ *P in mesocosms) produce a spectrum ranging from hydra effects to trophic cascades*

333 *for two-genotype populations.* In both simulations and the experiment, hosts and parasites are

334 added on days 0 and 28 (red tick mark), respectively. (a) With genotypes 1&2 present, the hydra

335 effect emerges given sufficient time as host density with parasites (solid) becomes higher than

336 without (dashed). (b) Mesocosms containing genotypes 1 & 2 experienced a hydra effect [mean

density across replicates with parasites (solid) or without (dashed), plotted at each time point; bars are standard error at each time point]. (c-f) With genotypes 1&3 or 2&3, a trophic cascade occurs in simulations and the mesocosm. (Parameters follow Table 1). For analyses, average mesocosm density was taken from day 48 to 76 (gray region, see Methods for mesocosm). Experimental time series shifted slightly horizontally for clarity. Compare simulations to Fig. 3's equilibrium outcomes and mesocosm time series to Fig. 5's mesocosm averages.

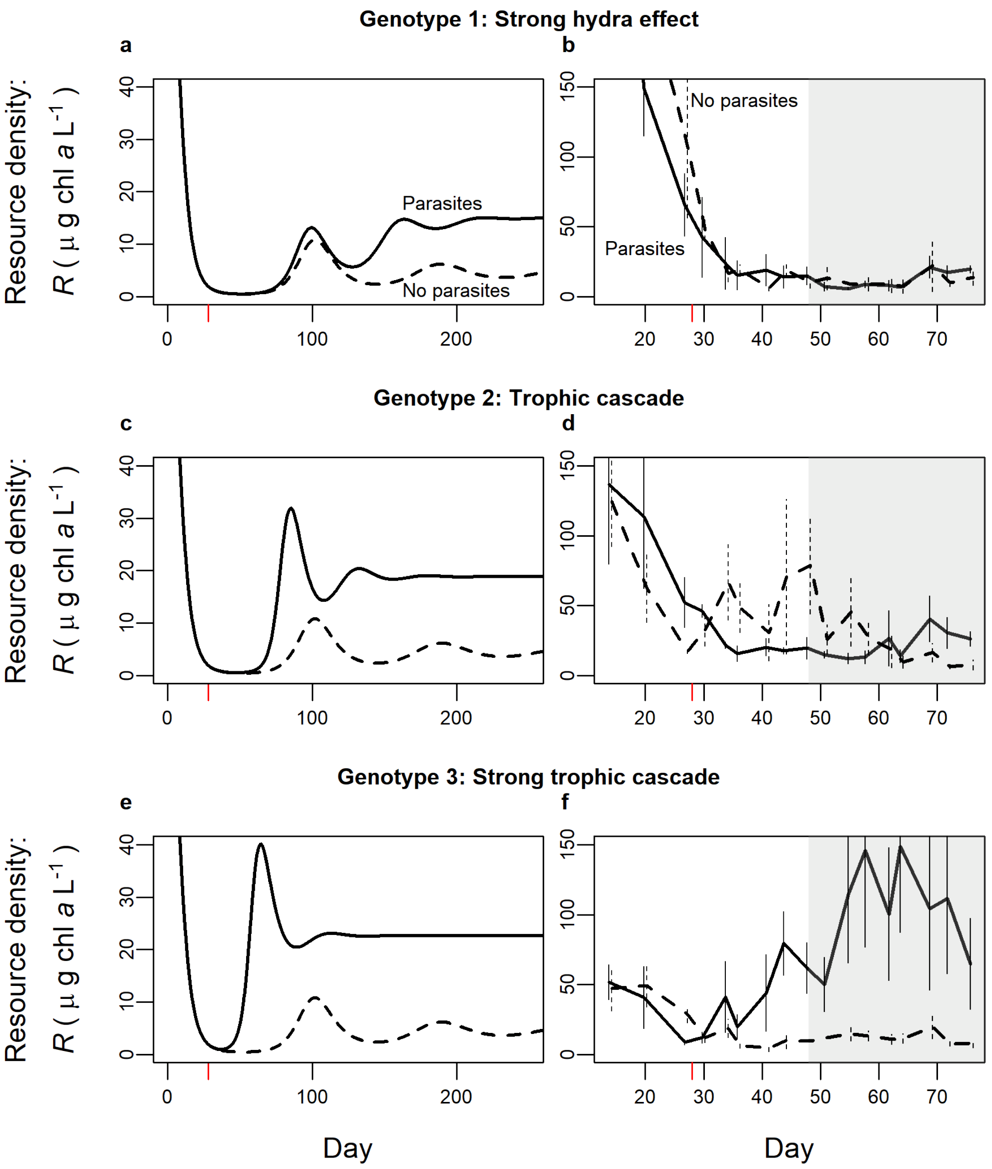

**Figure S5.** *Resource density in simulation and mesocosm time series for single-genotype treatments at high nutrients (*K *= 94.3 in simulations or 50 µg $L^{-1}$ P in mesocosms).* Resource release (one measure of cascade strength) compares resources with parasites (solid) to without (dashed). Treatments with stronger foraging depression and lower susceptibility experience smaller resource release in simulations (a) and mesocosms (b). Treatments with weaker depression and higher susceptibility (c-f) experience larger resource release, largely due to killing of hosts. For analyses, average mesocosm density was taken from day 48 to 76 (gray region, see Methods for mesocosm). Experimental time series shifted slightly horizontally for clarity. Compare simulations to Fig. 3's equilibrium outcomes and mesocosm time series to Fig. 5's mesocosm averages.

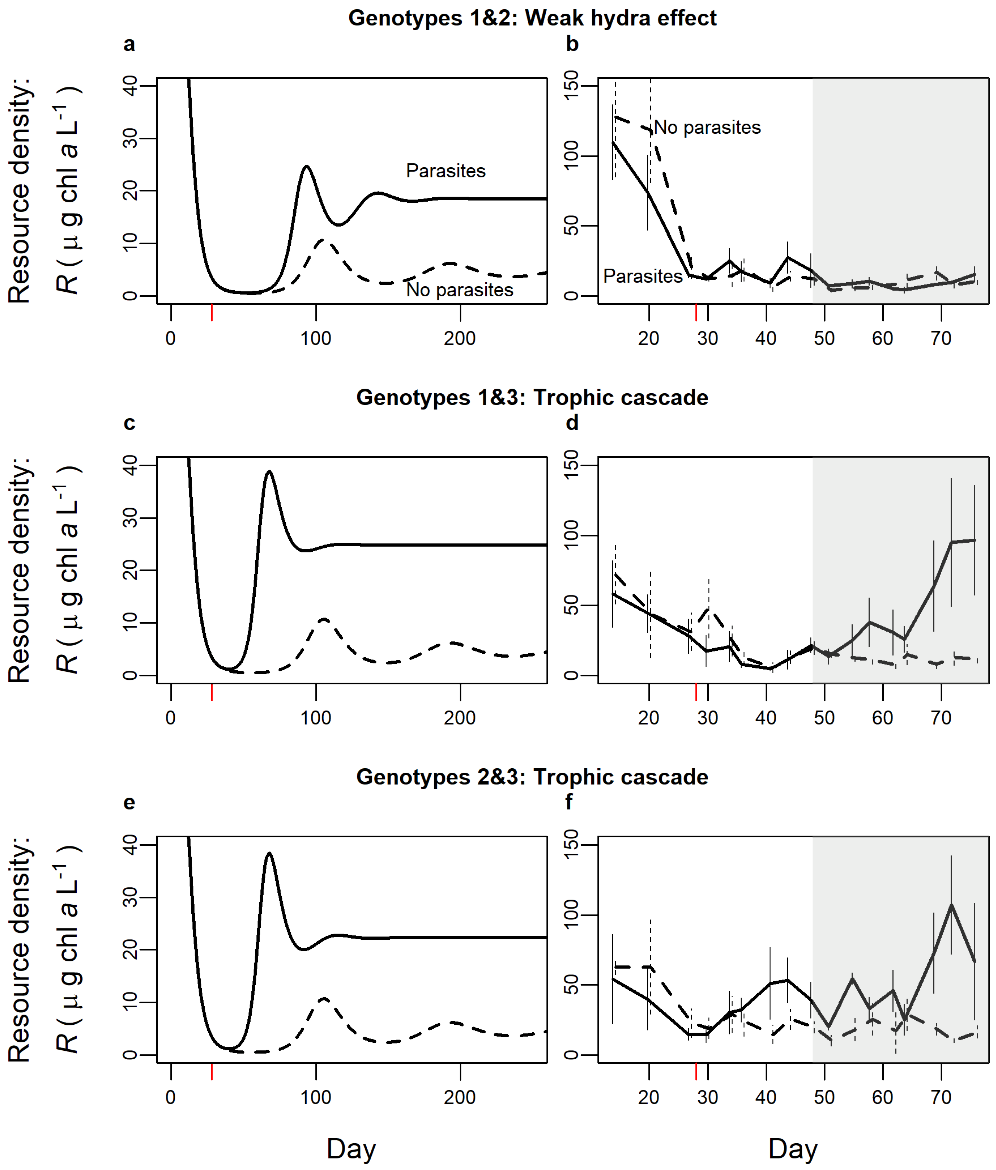

**Figure S6.** *Resource density in simulation and mesocosm time series for two-genotype treatments at high nutrients (*K *= 94.3 or 50 µg $L^{-1}$ P).* Resource release is measured by comparing resource density with parasites (solid) to without (dashed). For analyses, average mesocosm density was taken from day 48 to 76 (gray region, see Methods for mesocosm). Experimental time series shifted slightly horizontally for clarity.

***(h) Foraging depression can produce hydra effects in a model with direct transmission***

A model of direct transmission shows a parasite-driven hydra effect is not restricted to environmentally transmitted parasites. Here foraging depression functions in a similar manner, responding to the infectious stage (here $I$ instead of $Z$, so $\alpha$ has different units): $f(I) = f_0 e^{-\alpha I}$. This modification yields a three-dimensional model:

$$\frac{dR}{dt} = rR\left(1 - \frac{R}{K}\right) - f(I)(S + I)R \qquad \text{(S6a)}$$

$$\frac{dS}{dt} = cf(I)(S + I)R - dS - uf(I)SI \qquad \text{(S6b)}$$

$$\frac{dI}{dt} = uf(I)SI - (d + v)I \qquad (S6c)$$

Because this model is not our focus and our system does not provide biologically reasonable parameter values, we conduct a brief, mostly numerical analysis. The same analysis as for the mortality-only model case analytically proves that equilibrium host density ($H^*$) cannot be higher with disease than without if there is no foraging depression ($H^*_{Z+}/H^*_{Z-} < 1$ for $\alpha = 0$). Numerical analysis readily shows a hydra-like effect can arise when $\alpha > 0$ (e.g. for $c = 1.73$, $d = 0.0172$, $f_0 = 0.0368$, $K = 682$, $u = 0.994$, $v = 0.0193$, $w = 6.08$, $\alpha = 4.15 \times 10^{-6}$). In the direct transmission model, higher $v$ can increase host density but via a different mechanism. With environmental transmission, higher $v$ increases conversion of infected hosts into parasite propagules, $Z$ (leading to lower foraging rate). With direct transmission higher $v$ reduces density of infected hosts ($I$)

and reduces the spread of infection. Thus, with or without foraging depression, higher $v$ can lead to higher host density (Anderson 1979) by reducing disease (e.g. higher host density $c = 1$, $d = 0.05$, $f_0 = 0.03$, $K = 50$, $u = 1$, $w = 1$, $\alpha = 0$ and $v = 0.4$ than for $v = 0.3$). So, through a foraging depression mechanism, a model of direct transmission (eq. S6) produces similar hydra effects as that with environmental transmission (eq. 1). However, virulence mortality ($v$) can increase host density by a different mechanism in the two models.

### Section 2: Methodological information

#### *(a) Estimates of foraging depression (α) - Foraging rate assays – Fig. 1*

We estimated coefficients of foraging depression (α) with short-term assays of foraging rate. In these assays, we reared cohorts of individuals of a genotype until they reached five days old. Then, we separated them into 15 mL tubes receiving 1.0 mg mass/L *Ankistrodesmus falcatus* (~70 µg chl *a*/L). Foraging depression for genotypes 1 and 2 was measured in one experiment (Strauss *et al.* 2019) while those of genotype 3 were measured in a separate but similar experiment (unpublished until now). For genotypes 1 and 2, each tube received a dose of 0 ($N$ = 40 and 13, respectively), 75 (27 and 14), 200 (29 and 11), or 393 (38 and 14) parasite spores/mL. This high parasite density likely corresponds to densities in large epidemics in nature (Tara Stewart Merrill personal communication). A separate experiment for genotype 3 had different assay durations and spore treatments (but all else equal). These foraging rate assays lasted 2, 5, or 8 hours but these time differences did not affect estimated foraging rates so we combined them at 0 ($N$=36) and 400 spores/mL ($N$=36).

Despite these minor differences in design, both experiments then followed the same basic format. Control tubes interspersed through the experiment received the same treatment (i.e., algal density and spore dose) but without a zooplankton individual. All tubes were inverted approximately every 30 minutes while kept in the dark for up to 8 hours. At the end of the experiment, we removed animals, then measured remaining algae via *in vivo* fluorescence for control and treatment tubes with a Turner Trilogy Laboratory Fluorimeter. For each individual $S$ (1 host /15 mL), we determined foraging rate [$f(Z)$] from the algae remaining in the treatment tube ($R_f$) compared to the corresponding control tube ($R_0$) and the time lapsed, $t_E$ [i.e. $f = \ln(R_0/R_f)/(S\ t_E)$]. For a small number of tubes, algal concentration was higher for the treatment tube than the control tube, either due to death of the animal or a molting event (more likely). These pairings of treatment and control tube were eliminated from the analysis.

For each genotype, we then fit a model of foraging depression to the foraging rate data. Using the non-linear least-squares fitting function in R (R Core Team 2019), we fit foraging rate as a function of spores, $Z$ (following Strauss *et al.* 2019): $f(Z) = f_0 e^{-\alpha Z}$, where $f_0$ is the foraging rate without spores ($Z$=0), and $\alpha$ is the coefficient of foraging depression (Fig. 1a, used for model eq. 1). We found 95% confidence intervals for each genotype's $\alpha$ by bootstrapping ($10^4$ times). To bootstrap α for each genotype, we constructed sample datasets, retaining dataset size, by randomly sampling foraging rate-spore density pairs within genotype and with replacement. We then estimated confidence intervals from the distribution of $\alpha$ values fit to each bootstrapped dataset (following Efron & Tibshirani 1993).

#### *(b) Mesocosm experiment*

Each mesocosm was housed in a 75-liter acid washed polyethylene tank in a climate-controlled room held at approximately 21°C. We filled tanks to 50 L with 80% tap water (treated with Kordon Amquel Plus and Novaqua plus) and 20% filtered (Pall A/E: 1 μm) lake water.

Water loss from evaporation was replaced with further additions during the experiment. Low nutrient tanks received 5 μg $L^{-1}$ P (as $K_2HPO_4$) with corresponding nitrogen (as $NaNO_3$) while high nutrient tanks received 50 μg $L^{-1}$ P; N:P ratio was 20:1 by mass. Nutrients were replenished twice weekly throughout the experiment to account for an estimated (exponential) 5% per day loss rate. All tanks were inoculated with 2 mg (by dry weight) of the green alga *Ankistrodesmus falcatus* 7 days before hosts were introduced (algae on day -6, hosts on day 1) and allowed to grow on a 24 hr light cycle to reach a high enough algal density to support hosts.

We added hosts to each tank on day 1 (10 hosts $L^{-1}$); hosts then grew 27 days before addition of 4660 fungal spores $L^{-1}$ (day 28) to the disease treatment tanks. Isoclonal host lines obtained from Midwestern (MI, USA) lakes were cultured in the laboratory while spores (from Baker Lake, Barry Co, MI, USA) were cultured by passage through live hosts. Genotype 1 was 'Bristol 10', genotype 2 was 'A4-3', and genotype 3 was 'Standard'. Tanks received a 16 L: 8 D light cycle after host addition. Twice a week on days 14-86, we sampled 1 L of tank water, sieving animals through 80 μm mesh to destructively sample hosts. We visually counted and diagnosed hosts for infection using dissecting microscopes (40-50X).

***(c) Determining experimental densities – Figs 5, S7, S8***

Several outliers in the experimental mesocosm populations were removed from the analyses. The majority were removed due to extinction of the host population, most often at low nutrients. The following were removed due to extinction:

- genotype 2: 2 at low *K*, *Z*-; 1 at low *K, Z*+
- genotypes 2&3: 1 at low *K, Z*-; 1 at low *K*, *Z*+
- genotype 3: 1 at low *K*, *Z*+

- genotype 1: 1 at high *K*, *Z*-

Future modeling work may account for stochastic extinction to gain insight from these populations. Another population (genotypes 2&3, low *K*, *Z*-) was removed due to contamination with the focal fungal parasite at an unknown date. One population of genotype 1 with high nutrients and parasites present was removed due to extremely low host density, possibly due to chemical contamination. This population's Cook's distance was > 4X the mean for host density (corresponding to 95$^{th}$ percentile); such a deviation is uncharacteristically low for this genotype, even at lower nutrient supply (J. Walsman, personal observation).

Population averages were taken over a 28-day time window (days 48 – 76) to best estimate quantities relevant to our theoretical models. On day 48, most populations in disease treatments (inoculated day 28) began to display sufficient visible infections. We ended on day 76 based on visual inspections of mesocosms and previous experiments. Around this time, mesocosms accumulate detritus and dynamics become less consistent across replicates. Then, over this 28-day window, we calculated averages as area under the curve divided by time. These averages provide closest comparison to model equilibria (eq. 2).

***(d) Nutrients and susceptibility increase prevalence in experimental populations – Fig S7***

The model predicts that increased resource carrying capacity (*K*) and host susceptibility (*u*) increase prevalence. This pattern holds with foraging depression in the model (see results of numeric search in Appendix section 1) and only grows stronger with a negative correlation between *u* and foraging depression (see Fig. 4). We tested the statistical effects of nutrients and susceptibility on prevalence with a beta regression. While a linear model finds the same qualitative result, a beta regression is better suited for prevalence, which is bounded between zero and one (Ferrari & Cribari-Neto 2004; Mangiafico 2016). We implemented the beta regression using the betareg package in Rstudio (R Core Team 2019) and the default "logit" link function. Diagnostic plots (following Ferrari & Cribari-Neto 2004) supported the use of the beta regression. The regression indicated that susceptibility (P = 0.0067) and nutrients (one-sided P-value = 0.0198) both increased prevalence.

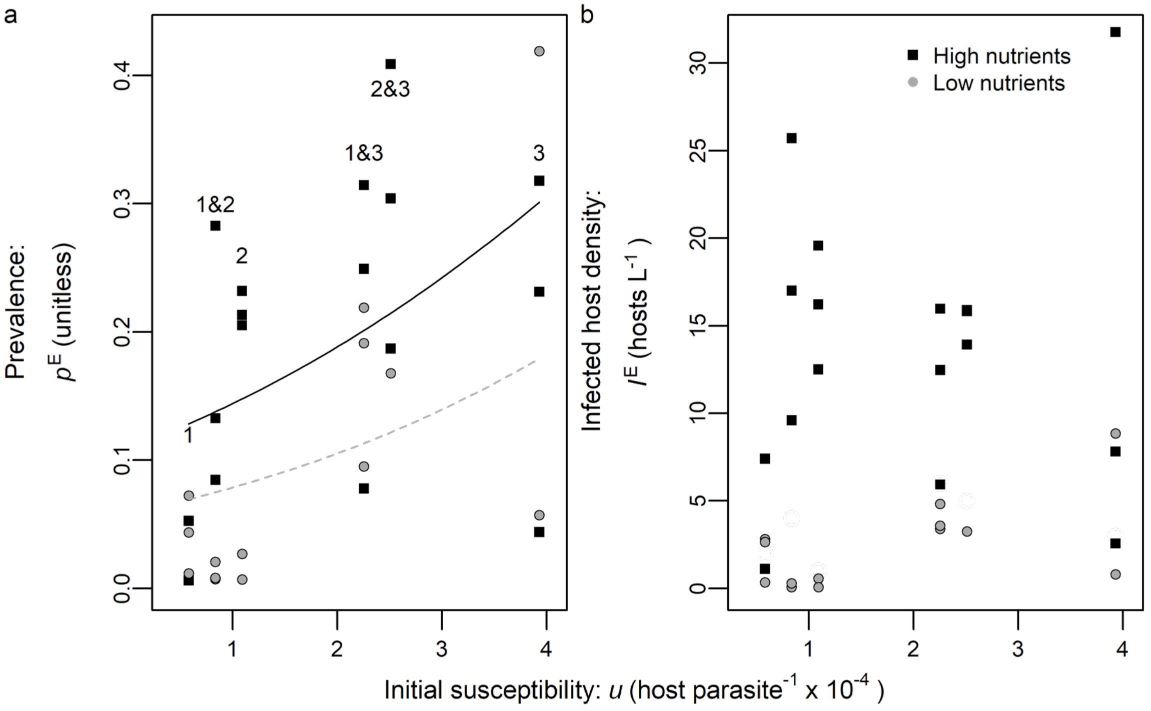

**Figure S7.** *Prevalence of infection and density of infected hosts in mesocosms.* Each point is a mesocosm population averaged over time. Gray circles: low nutrients; black squares: high nutrients. Vertical groupings are genotype treatments (with number labels on top). With increasing initial susceptibility ($u$) and nutrients ($K$), (a) prevalence ($p^E$) increases. (b) Infected host density ($I^E$) can reach high density compared to initial infection dose (~equivalent to two infected hosts per 50 L, or 0.04 $L^{-1}$). This increase in infection density demonstrates that parasite epidemics were self-sustaining in experimental populations.

Experimental mesocosms experienced self-sustaining, multi-generational epidemics of varying sizes. In some populations, especially those with high susceptibility to infection (see beta regression), nearly half of the population became infected (Fig. S7a). Epidemics were initialized with 4,660 spores/L or 233,000 spores/population. This is roughly equivalent to the spores released from two heavily infected hosts per population. Many disease populations attain a density of infected hosts greater than 10/L (see Fig. S7b). With 50 L populations, this corresponds to more than 500 infected hosts/population. Thus, initial infections (of animals reared to produce spores used to inoculate mesocosms) resulted in secondary and (very likely) tertiary infections, creating self-sustaining parasite epidemics. These self-sustaining parasite epidemics distinguish our experiment from many laboratory experiments with one parasite generation and/or donor-controlled parasite abundance. These less dynamic parasite populations are tractable and useful for reducing experimental variation. But self-sustaining epidemics over multiple host generations match assumptions of our dynamic model of feedbacks between interacting populations of parasites, hosts, and resources. Perhaps more importantly, these dynamic feedbacks more closely resemble those during epidemics in nature. Furthermore, such feedbacks (especially for resources) are required to produce the hydra effect.

Infected host density can also help approximate parasite propagule densities. The model predicts equilibrium parasite density as $Z^* = \sigma(d+v)I^*/m$ (from eq. 1d). Given reasonable parameter values (see Table 1), 30 infected hosts/L (population average of upper right point in Fig. S7b) corresponds roughly to 1.15 x $10^5$ spores/L. The highest transient infected host density observed in any treatment in the epidemic window was 82 infected hosts/L. Assuming the conversion is still roughly appropriate, this corresponds to 3.14 x $10^5$ spores/L. Thus, the span of spore doses used in the foraging depression assay (see Fig. 1) likely corresponds to the range of parasite propagule densities host experienced in the mesocosms.

***(e) Mapping model results onto predictions of main effects and interactions – Table S1***

Whether or not foraging depression is present in the model, the model predicts the same main effects and interactions for disease, nutrients, and susceptibility. Higher susceptibility should always strengthen resource release and host suppression (Figs 4c, d). Higher carrying capacity of the resource should always strengthen resource release (Fig. 4a) and may strengthen or weaken host suppression (only weaken shown in Figs 4b). Given trait measurements, disease should usually decrease host density and should always increase resource density (see Figs 3, 4) and higher carrying capacity should weaken host suppression (Fig. 4a, b).

We tested the effects of disease and its interactions with susceptibility and nutrients (resource carrying capacity) on experimental resource and host density. Each data point is the average for a given population in a unique mesocosm, ensuring independence of observations. We fit a linear model (eq. S7) to $\log_{10}$ resource and host density in Rstudio (R Core Team 2019). Diagnostic plots supported the assumptions of linearity, homoscedasticity, and normally distributed error.

We found the effects of treatments on experimental resource ($R^E$) and host ($H^E$) densities using linear model fits to $\log_{10}$ mesocosm densities. The models take the following form:

$$\log_{10}(R^E) = r_0 + r_1K + r_2u + r_3Z + r_{13}KZ + r_{23}uZ + \varepsilon_R \quad \text{(S7a)}$$

$$\log_{10}(H^E) = h_0 + h_1K + h_2u + h_3Z + h_{13}KZ + h_{23}uZ + \varepsilon_H \quad \text{(S7b)}$$

We modeled $\log_{10}$ resource ($R^E$; eq. S7a) or host ($H^E$; eq. S7b) density in the experiment as a function, from left to right, of an intercept ($r_0$ and $h_0$), nutrients (represented by carrying capacity, $K$), susceptibility ($u$), and disease ($Z$) with $K$ x $Z$ and $u$ x $Z$ interactions and an error term ($\varepsilon_R$ or $\varepsilon_H$ following a Gaussian distribution). To aid interpretation of regression coefficients (i.e., the $r_j$ and $h_j$ parameters for resources and hosts, respectively), we centered the independent numerical variables ($K$ and $u$) to have mean zero. Thus, we used values above or below the mean (zero) to predict the variable's effect on density. Then, for the categorical disease variable, we used a coding scheme that more naturally matched predictions of the differential equation model (eq. 1). That model did not predict meaningful overall effects of $u$ on density; hence, we did not code disease a more traditional way (which would fit main effects of $K$ and $u$ to the data overall, i.e. $Z$- = -1 and $Z$+ = 1). Instead, we coded the categorical disease treatment so that no disease ($Z$-) is 0 and disease ($Z$+) is 1. Therefore, no disease is the default in this linear model; the main effects of $K$ and $u$ are provided without disease. This choice, then, allowed comparison to clear predictions.

Mapping theory predictions onto fitted coefficients is straight-forward because the derivative of equilibrium density (on an arithmetic scale) has the same sign as the derivative of $\log_{10}$ density. For example, higher carrying capacity ($K$, related to increased nutrients for algae) increases equilibrium host density in the absence of parasites (d/d$K$ $H^*_{Z-}$ > 0; derived from eq. 2c). Thus, $K$ must also increase $\log_{10}$ host density [d/d$K$ $H^E_{Z-}$ > 0 implies d/d$K$ $\log_{10}(H^E_{Z-}) = h_1 >$ 0; eq. S7b]. The same logic predicts the signs of other regression coefficients: $r_1 = 0$, $r_2 = 0$, $h_2 =$ 0 (see theory predictions and experimental outcomes compared in Table S1).

Similarly, the model predicts main effects of the categorical disease treatment and its interactions. The model predicts disease will increase resources ($R^*_{Z+} > R^*_{Z-}$; $r_3 > 0$) and usually decrease host density ($H^*_{Z+} < H^*_{Z-}$; $h_3 > 0$). Interaction effects are predicted by how a variable (carrying capacity $K$ or susceptibility $u$) influences density ratio. For example, host density ratio decreases with susceptibility in the model [d/d$u$ ($H^*_{Z+}/H^*_{Z-}$) < 0 because $H^*_{Z+}$ decreases with $u$, Fig. 4d]. Thus, $\log_{10}(H^*_{Z+}/H^*_{Z-})$ also decreases with $u$. A negative effect of $u$ on $\log_{10}$ host ratio equates to $h_{23} < 0$. Thus, there should be a negative interaction between susceptibility and disease for experimental host density because the model predicts parasite-driven host suppression is stronger when hosts are more susceptible.

The model predicts most features of resource release in the experiment. The addition of disease significantly increased resources (Fig. S8a, b). Susceptibility to infection ($u$) had no impact on algal density without disease (flat gray line Fig. S8b). But, as predicted, resource release was magnified by higher susceptibility (positive $u$ x $Z+$ interaction; black slope higher than gray in Fig. S8b and increasing resource ratio in Fig. 5a). All of these results were consistent with predictions (Table S1). However, a few small inconsistencies between model and experiment also arose. The effect of nutrient supply on resources differed somewhat from that predicted by the model equilibrium, likely due to transient dynamics. In the model at equilibrium and without disease, hosts graze resources down to their minimum resource requirement, which does not depend on resource carrying capacity (eq. 2a). Before reaching equilibrium, transient resources can increase with carrying capacity until hosts depress resources to the hosts' minimal requirement. This likely explains why, in the mesocosms, algal resources in the absence of disease ($R^E_{Z-}$) increased somewhat with nutrient supply (significant positive effect of $K$; see Fig. S8a, Table S1); increasing $R^E_{Z-}$ with nutrient supply likely also weakened the $K$ x $Z$ interaction for resources. Thus, the treatment effects on algal density were largely (but not entirely) consistent with the model.

The model largely predicts drivers of host suppression as well. Higher nutrient supply ($K$) should and did increase host density ($H^E_{Z-}$) without disease (Fig. S8c, Table S1). Counter to the model, higher susceptibility (*u*) did increase host density without disease (positive gray slope Fig. S8d). This relationship might have arisen due to differences in non-focal traits of these genotypes (Strauss et al. 2015). Nonetheless, parasites suppressed host density (Figs S8c, d). Host suppression weakened non-significantly with higher nutrient supply (i.e., non-significant, positive *K* x *Z*+ interaction for host density). Higher susceptibility, as predicted, did amplify host suppression (negative *u* x *Z*+ interaction, as predicted; black line [*Z*+] had lower slope than gray line [*Z*-] in Fig. S8d and decreasing host ratio in Fig. 5b). So, susceptibility strengthened host suppression while nutrient supply did not significantly affect it (Fig. 5b).

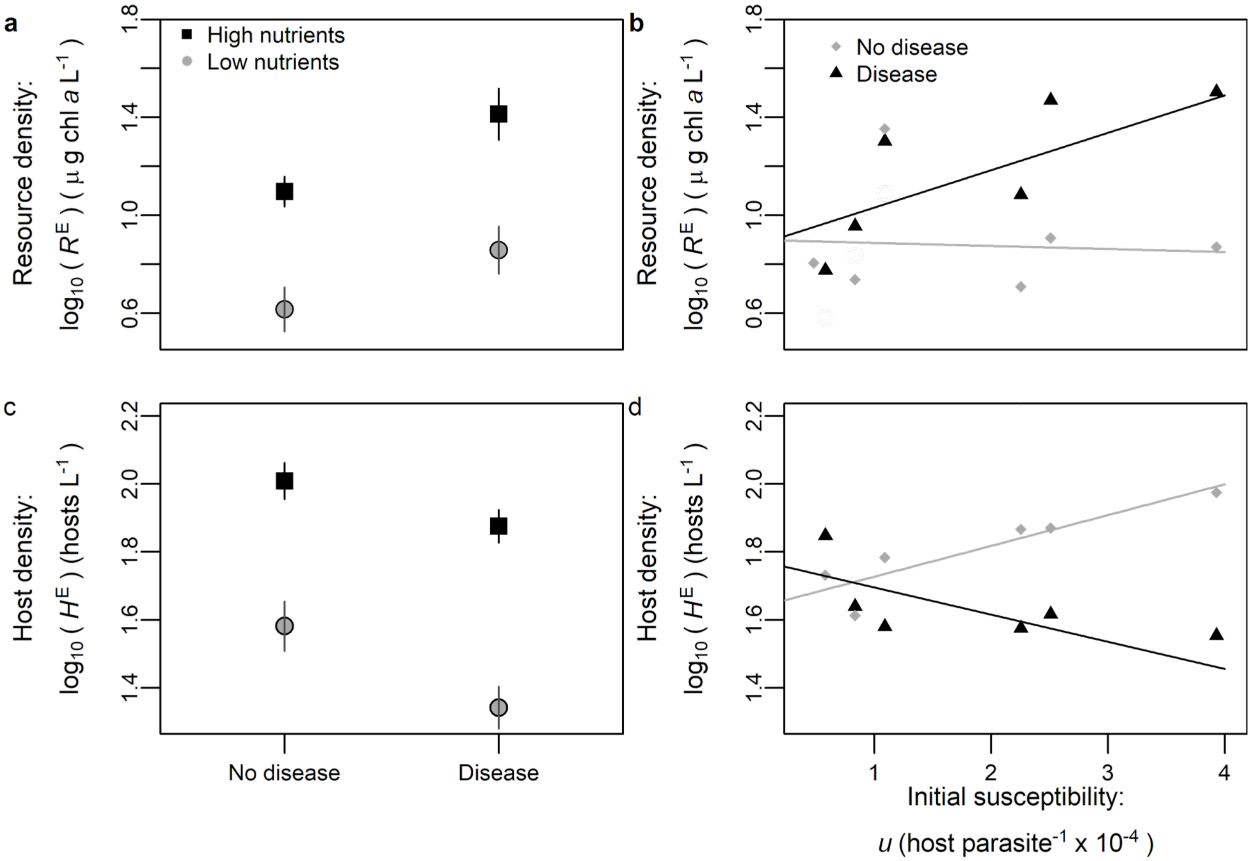

**Figure S8.** *Parasites drive cascades modulated by susceptibility more than carrying capacity.*

Each point represents an average over time and populations within a group of treatments of $\log_{10}$ density. First column is grouped by nutrient supply and disease treatment. Nutrient supply treatments: high (black squares) and low (gray circles; see text). Susceptibility (*u*) was manipulated using one of three single-genotypes (1, 2, or 3) or (initially) a 50:50 mixture of two (e.g. 1&2), creating a range of *u*. *Algal resources*: (a) Nutrient supply (*K*) and disease (*Z*+) increase experimental resource density ($R^E$; compare Fig. S1c) with no interaction (see Table S1). Second column is grouped by susceptibility and disease treatment. (b) Susceptibility does not affect resource density without disease (diamonds along flat gray line) but increases resources with disease (triangles increasing with black line; as in Fig. S1c; *u* x *Z*+ interaction). *Plankton hosts:* (d) Nutrient supply increases and disease decreases host density ($H^E$; as in Fig. S1e) with no interaction. (e) Susceptibility increases host density without disease (diamonds along gray line) but decreases it in the presence of disease (triangles along black line; compare Fig. S1e; *u* x *Z*+ interaction).

**Table S1.** *The model (eqs. 1, 2) largely predicts GLM treatment effects in a mesocosm experiment*. Theoretically predicted or experimentally determined effects of nutrient supply (*K*), host susceptibility (*u*), or disease (*Z*+) treatments (Trmt) on resource (*R*) or host (*H*) density. Here, disease-free (*Z*-) treatments are the default. Hence, main effects of *K* and *u* denote their effects without disease. A '+/-' means that theory predicts a potential positive or negative effect while NS denotes non-significant results (e.g. "NS+" is a non-significant positive trend). Theoretical predictions are drawn from equilibrium densities (eq. 2) mapped onto GLM coefficients (see Appendix: Section 2). Many of the predictions are general but some depend on biologically relevant parameter values. Experimental ('E') values are parameter estimates ($r_i$, $h_i$) from a linear model (eq. S3) predicting $\log_{10}$ mean experimental density ($R^E$ and $H^E$) averaged over time and treatment. P-values are provided.

| **Trmt** | **Resources (*R*)** | | | **Hosts (*H*)** | | |
|---|---|---|---|---|---|---|
| | **Theory $R^*$** | **Exp. $R^E$** | **P-value** | **Theory $H^*$** | **Exp. $H^E$** | **P-value** |
| ***K*** | d/d$K$ $R^*_{Z-}$: **0** | 0.011 | 0.0002 | d/d$K$ $H^*_{Z-}$: + | 0.00946 | $6.28 \times 10^{-7}$ |
| ***u*** | d/d$u$ $R^*_{Z-}$: **0** | -0.0124 | 0.806 | d/d$u$ $H^*_{Z-}$: **0** | 0.0906 | 0.00612 |
| ***Z*+** | $R^*_{Z+} > R^*_{Z-}$ | 0.29 | 0.00126 | $H^*_{Z+} < H^*_{Z-}$ | -0.182 | 0.00137 |
| ***K* x *Z*+** | d/d$K$ $R^*_{Z+}/R^*_{Z-}$: + | 0.000731 | 0.841 | d/d$K$ $H^*_{Z+}/H^*_{Z-}$: **+/-** | 0.00286 | 0.241 |
| ***u* x *Z*+** | d/d$u$ $R^*_{Z+}/R^*_{Z-}$: + | 0.0731 | 0.0277 | d/d$u$ $H^*_{Z+}/H^*_{Z-}$: **-** | -0.171 | 0.000503 |

***(f) Statistical significance of experimental hydra effects – Fig. 5***

Host density appeared to be higher with disease than without disease (a hydra effect) for treatments with high nutrient supply and with host genotype 1 or genotypes 1 and 2 combined. Necessary removal of outlier populations (see above: Determining experimental densities) and the occurrence of a hydra effect for only certain genotypes provided a small number of replicate populations (2 with parasites and 2 without for genotype 1; 3 with parasites and 3 without for genotypes 1&2). Nine repeated measurements of each population over time provide additional statistical power. But these repeated measurements are autocorrelated. To account for this autocorrelation, we used a nested ANOVA, with time nested within individual mesocosm and individual mesocosm nested within disease treatment. We performed the nested ANOVAs in R

(R Core Team 2019) with host density and log host density. Host density (whether or not it was log transformed) was significantly higher with disease for both genotype treatments. However, the homoscedasticity assumption of nested ANOVAs, as diagnosed with a residuals vs fitted plot, was satisfied better by log-transformed host density. Normal Q-Q plots also revealed residuals of log-transformed host density to be approximately normal for both genotype treatments. Thus, we report results for log host density. As reported in the text, host density was significantly higher with disease than without disease for genotype 1 alone ($P = 0.00748$) as well as the mixed genotype 1 and 2 treatment ($P = 0.0201$).

### References

Anderson, R.M. (1979) Parasite pathogenicity and the depression of host population equilibria. *Nature,* **279,** 150-152.

Ebert, D., Joachim Carius, H., Little, T. & Decaestecker, E. (2004) The evolution of virulence when parasites cause host castration and gigantism. *American Naturalist,* **164,** S19-S32.

Ebert, D., Lipsitch, M. & Mangin, K.L. (2000) The effect of parasites on host population density and extinction: Experimental epidemiology with Daphnia and six microparasites. *American Naturalist,* **156,** 459-477.

Efron, B. & Tibshirani, R.J. (1993) *An Introduction to the Bootstrap*. Chapman & Hall, New York.

Ferrari, S. & Cribari-Neto, F. (2004) Beta regression for modelling rates and proportions. *Journal of applied statistics,* **31,** 799-815.

Hall, S.R., Becker, C. & Cáceres, C.E. (2007) Parasitic castration: a perspective from a model of dynamic energy budgets. *Integrative and Comparative Biology,* **47,** 295-309.

Mangiafico, S.S. (2016) *Summary and Analysis of Extension Program Evaluation in R, version 1.18.1.*

McKay, M.D., Beckman, R.J. & Conover, W.J. (2000) A comparison of three methods for selecting values of input variables in the analysis of output from a computer code. *Technometrics,* **42,** 55-61.

R Core Team (2019) R: A language and environment for statistical computing. R Foundation for Statistical Computing, Vienna, Austria.

Shurin, J.B., Borer, E.T., Seabloom, E.W., Anderson, K., Blanchette, C.A., Broitman, B., Cooper, S.D. & Halpern, B.S. (2002) A cross-ecosystem comparison of the strength of trophic cascades. *Ecology Letters,* **5,** 785-791.

Shurin, J.B. & Seabloom, E.W. (2005) The strength of trophic cascades across ecosystems: predictions from allometry and energetics. *Journal of Animal Ecology,* **74,** 1029-1038.

Strauss, A.T., Hite, J.L., Civitello, D.J., Shocket, M.S., Cáceres, C.E. & Hall, S.R. (2019) Genotypic variation in parasite avoidance behaviour and other mechanistic, nonlinear components of transmission. *Proceedings of the Royal Society B,* **286,** 20192164.
